# Divergent roles of DNA methylation, TRIM28, and p53 surveillance in human embryonic and trophoblast stem cells

**DOI:** 10.1101/2025.10.19.683051

**Authors:** Deepak Saini, Megan S. Katz, Skye G.A. Beck, Taylor E. Ireland, Soma Ray, Asef Jawad Niloy, Radman Mazloomnejad, Anna Fleming, Sophia C. Vetrici, Jessica K. Cinkornpumin, Tanja Sack, Stephanie Duhamel, Rima Slim, Morag Park, Jennifer M. Frost, Soumen Paul, Raquel Cuella Martin, William A. Pastor

## Abstract

Transcriptional regulation of transposons and genes by TRIM28 and 5mC is critical for proper mammalian embryonic development, but the specific roles for these mediators in human embryonic and placental lineage remain unclear. We find that loss of TRIM28 has a limited effect on global transposon expression and instead results in upregulation of genes proximal to TRIM28-bound Long Terminal Repeats (LTRs) in both human trophoblast stem cells (hTSCs) and human embryonic stem cells (hESCs). MER11A elements show especially strong regulatory importance in hTSCs: these elements are bound by both TRIM28 and placental transcription factors and show both heterochromatic and euchromatic features. Some genes are positively regulated by MER11A elements in hTSC basal state, while other MER11A-proximal genes show upregulation only upon TRIM28 deletion. By contrast, loss of DNA methylation in hESCs or hTSCs leads to a global increase in transposon expression. While many genic 5mC targets are shared in hESCs and hTSCs, we also observe evidence that a handful of genes important for somatic development are repressed by 5mC in trophoblast, while a small parallel set of placental genes are repressed by methylation in embryonic tissue. Interestingly, loss of DNMT1 causes hESCs to be rapidly lost from culture in a *TP53* and mitotic surveillance checkpoint-dependent manner, while hTSCs show little p53 response to DNMT1 loss or DNA damage generally, instead showing gradual mitotic defects and aneuploidy and slow loss from culture. This discrepancy may explain the higher frequency of karyotypic abnormality found in human placental cells. Together, this study charts the role of TRIM28 and DNA methylation in regulating embryonic and placental transcription and demonstrates divergent p53-dependent responses to genomic instability.

## INTRODUCTION

Methylation of the 5-position of cytosine (5mC) is a critical mechanism for gene silencing in many organisms. During mammalian development, most 5mC is lost during the first cycles of cell division, such that global levels of 5mC are quite low in the morula and pre-implantation blastocyst^1,2^. In the subsequent peri-implantation period, global patterning of 5mC occurs^1,3^. By this stage, the epiblast (embryonic) and trophoblast (placental) lineages have already diverged and acquire dramatically different genomic patterns of 5mC^3–5^.

Much remains unknown about the function of 5mC during human development, particularly in placenta. In murine embryonic and placental tissue, loss of key DNA methyltransferases (DNMTs) causes upregulation of genes involved in germ cell development (germline genes) as well as transposons^6,7^. During the pre-implantation window, when global 5mC levels are low, the repressive TRIM28 complex is critical for transposon silencing. *Trim28^-/-^* mice undergo embryonic lethality by E5.5^8^. Upon genome-wide methylation, 5mC supplants TRIM28 as the predominant transposon silencer^7,9^. The global 5mC level in the placenta is far lower than in the embryo^10,11^, raising the possibility that TRIM28 retains a significant role in transposon repression even after methylation patterning. It has also been observed that transposon sequences have been co-opted to function as enhancers in the placental lineage^12,13^, potentially allowing shifts in overall transcription that facilitate the rapid evolution of the placenta. Thus, another possibility is that transposons are simply less repressed in trophoblast.

There is strong evidence implicating imprinted genes, including genes selectively imprinted in placenta, in regulation of placental development^14,15^. Mouse embryos with maternal deficiency of *de novo* DNA methyltransferases, causing loss of maternal imprints, feature developmental abnormalities caused in substantial part by dysregulation of the imprinted gene *Ascl2*^16^. However, *ASCL2* is not imprinted in humans^17^. Errors of meiosis or fertilization leading to an androgenetic human embryo, in which maternal DNA and thus maternal imprints are absent, results in an abnormal conceptus called a hydatidiform mole^18^. Lost expression of the placentally imprinted tumour suppressor *CDKN1C* in hydatidiform moles results in loss of contact inhibition and overgrowth of cytotrophoblasts^19^. However, in addition to imprinted genes, hundreds of non-imprinted gene promoters are selectively methylated in the embryonic or placental lineage^20^. Whether these genes are repressed by 5mC is unclear; just because a gene has a methylated promoter does not mean that loss of 5mC will reactivate it. Neither is it known whether the repression of these genes has any biological importance.

Even the importance of 5mC in placental cell survival is unclear. In mice, embryonic stem cells (ESCs) and trophoblast-derived trophoblast stem cells (TSCs) can survive with mutations of all three DNA methyltransferases and thus no DNA methylation^21,22^. *Dnmt1^-/-^ Dnmt3a^-/-^ Dnmt3b^-/-^* murine ESCs (mESCs) can self-renew normally but cannot differentiate or contribute to chimeras^21,22^. *Dnmt1^-/-^ Dnmt3a^-/-^ Dnmt3b^-/-^* mTSCs can likewise self-renew, albeit with upregulation of some differentiation genes, and even show some contribution to placenta in chimera assays^22^. Human ESCs (hESCs) by contrast die rapidly upon loss of the maintenance methyltransferase DNMT1^23^; the importance of 5mC in human trophoblast cell viability is unknown.

Here we determine the role of 5mC in gene and transposon regulation in cells of human embryonic and placental origin, and the cellular response to loss of methylation. We find that TRIM28 primarily acts by repressing LTR elements from acting as enhancers, while DNA methylation suppresses a wide range of transposons as well as select genes specific to embryonic or placental lineage. Finally, we observe a muted p53 response in hTSCs in response to DNA damage.

## MATERIALS AND METHODS

### Cell culture of human trophoblast stem cells and human embryonic stem cells

All human trophoblast stem cells used in this study were procured from the Arima lab (Tohoku Graduate school of Medicine, Japan). Wild-type CT27 (female), CT29 (male) and CT30 (female) cells originate from first trimester placenta^24^. hTSCs were cultured on 2.5 μg/mL laminin-511 (Millipore-Sigma, CC160-1050UG) pre-coated plates and media was refreshed every two days. Cells were passaged when up to 90% confluent using 30% TrypLE (Gibco^TM^, 12604021) and passaged at ratios between 1:5 or 1:10. hTSCs were maintained and cultured in media composing of DMEM-F/12 (Gibco^TM^, 11320033) media supplemented with 0.3% BSA (Millipore Sigma, A9205-50ML), 1% Insulin-Transferrin-Selenium-Ethanolamine (Gibco^TM^, 51500056), 1% Penicillin/Streptomycin (Gibco^TM^, 15140163), 0.1 mM 2-mercaptoethanol (Thermo Scientific, 21985023), 50 ng/mL EGF (StemCell™, 78006.2), ES-qualified fetal bovine serum (Gibco^TM^, 20439024), 1.5 μg/mL L-ascorbic acid, 2mM CHIR99021 (Cayman Chemical Company, 13122), 0.5 mM A83-01 (Selleck Chemicals, S7692), 1 mM SB431542 (Cayman Chemical Company, 13031), 5 μM Y-27632 (Cayman Chemical Company, 10005583) and 0.8 mM Valproic Acid (Millipore Sigma, p6273-100ML).

hTSCs with differentiated to 3D-syncytiotrophoblasts (3D-STB) using a published protocol^25^. hTSCs were first dissociated with TrypLE and 3x10^5^ cells/well were transferred to a non-adherent 6-well plate in 3mL of following media conditions: DMEM-F/12 supplemented with 0.3% BSA, 1% Penicillin/Streptomycin, 1% Insulin-Transferrin-Selenium-Ethanolamine, 0.1 mM 2-mercaptoethanol, 2.5 μM Y-27632, 4% Knockout™ Serum Replacement (Gibco^TM^, 10828028), 5 μM Forskolin (Cayman Chemical Company, 11018), 50 ng/mL EGF. On day 2 of culture, an additional 3 mL of media was added to each well and cells were collected on day 4 for analysis.

H9 hESC (female) used in this study were procured from Wicell. hESC were cultured on hESC-qualified Matrigel matrix (Corning^®^, CLS354277) pre-coated plates and media was refreshed every day. Cells were passaged when up to 90% confluent using Gentle Cell Dissociation Reagent (StemCell^TM^, 100-0485) and passaged at ratios between 1:20 to 1:50. H9 hESCs were then maintained and cultured using mTESR^TM^ plus media (StemCell^TM^, 100-1130). In experiments that involve passaging cells to single cells, Y-27632 was supplemented to enhance survival.

### Genetic ablation of stem cell lines

*DNMT1^exon33/exon33^, TP53^Exon5/Exon6^, TP53BP1^Exon4/Exon4^, USP28^Exon4/Exon4^ and TRIM28^Exon4/Exon4^* hTSC and hESC lines were generated using the sgRNA targeting flanking exons of interest. SpCas9-2NLS and sgRNA (Synthego) were delivered using the Lonza Biosciences 4D-Nucleofector to electroporate ribonuclear particles into human stem cells. For each experiment 300,000 cells were prepared according to the manufacturer’s protocols and were nucleofected using the CA-137 protocol. Cells were then hastily seeded to pre-warmed media after nucleofection, and media was changed the morning after. To generate stable clonal mutant lines, cells were passaged onto a 6-well plate at low density (approximately 2000 cells per 9.6 cm^2^ well) and were colony picked and expanded. Individual clones were validated using PCR, immunofluorescence microscopy or western blot. Control lines were generated by nucleofection with a non-targeting sgRNA and were treated identically unless otherwise indicated.

The following sgRNA sequences were used:

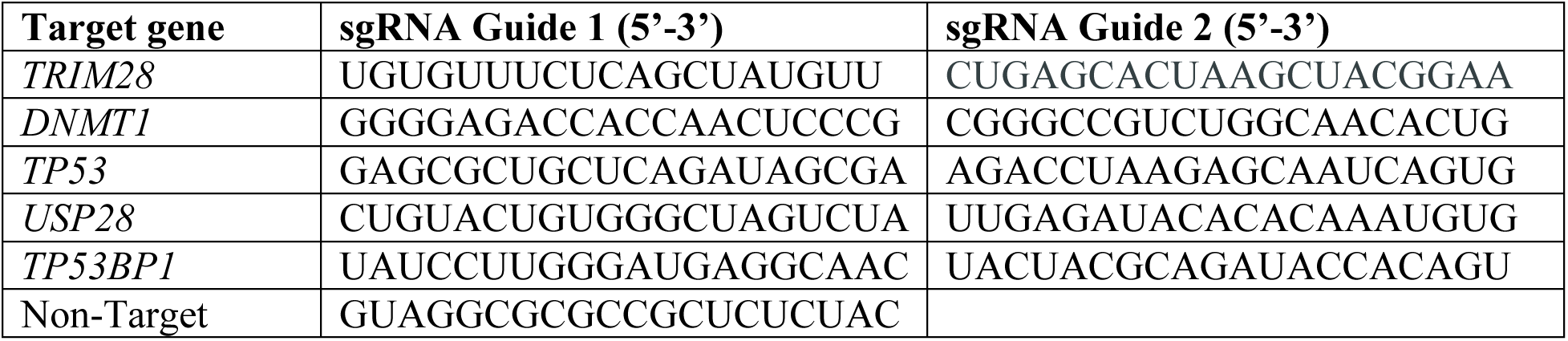

### Generation of CRISPR interference (CRISPRi) plasmids and MER11A CRISPRi CT30 hTSC cell lines

CRISPRi sgRNA targeting MER11A proximal to the LEP gene were designed through the CRISPick software (PMID: 29431740, 32661438). sgRNA was then cloned into the following CRISPRi plasmid backbone, pLV hU6-sgRNA hUbC-dCas9-KRAB-T2a-Puro (Addgene, 71236) (PMID: 26501517) using BsmBI-v2 restriction enzyme (New England Biolabs, R073S). CRISPRi plasmids were verified using sanger sequencing. Viral constructs containing the CRISPRi plasmids of interest were generated using HEK293Ts. HEK293T cells were plated in a 6-well plate and maintained in DMEM-F/12 supplemented with 10% HyClone™ Bovine Calf Serum (Cytvia, SH3007203) and 1% penicillin-streptomycin until 70% confluency. HEK293T cells were then used for lentiviral production using Lipofectamine 3000 (Life Technologies, L3000008) using manufacturer’ instructions and the following plasmids: 1250 ng cloned CRISPRi plasmids (Addgene, 71236), 625 ng psPAX2 (Addgene, 12260) and 625 ng pMDG.2 (Addgene, 12259). HEK293T media was changed the following morning to hTSC media and then collected after 24 hours. The viral supernatant was filtered through a 0.45 μm PES filter and frozen down at -80 °C. hTSCs were seeded in a well of a 24-well plate and the following morning media was replaced with 50% viral supernatant and 50% hTSC media supplemented with 10 µg/mL polybrene. After 24 hours, media was washed with PBS and replaced with 100% hTSC media. 3-4 days after viral infection, hTSC media was supplemented with 1 µg/mL puromycin to select for hTSCs containing the CRISPRi construct. Control empty vector hTSC were generated along side the MER11A CRISPRi mutants and were treated identically unless otherwise indicated.

For the following primers were used for CRISPRi sgRNA targeting:

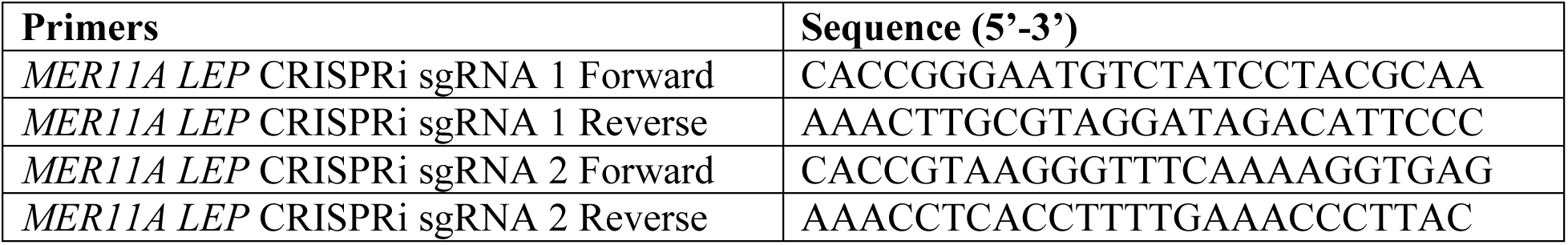

### Transdifferentiation of *TP53^-/-^* H9 hESC

H9 human embryonic stem cells were also transdifferentiated to trophoblast stem cells. hESCs were first reverted to a naïve state following an established 5iLAF protocol^26^. 5iLAF media was composed of: 50% Neurobasal (Gibco™, 21103049) and 50% DMEM-F/12, supplemented with 1x N-2 supplement (Gibco™, 17502001), 1x B-27 (Gibco™, 17504044), 2 mM GlutaMAX (Gibco™, 35050061), 1x MEM non-essential amino acids (Gibco™, 11140076), 1% penicillin-streptomycin, 50 mg/mL bovine serum albumin (Wisent, 809-098-EL), 0.1 mM 2-mercaptoethanol, 1 μM PD032901 (Axon, 1408), 1 μM IM-12 (Axon, 2511), 0.5 μM SB590885 (Axon, 2504), 1 μM WH-4023 (Axon, 2381) and 10 μM Y-27634 (Axon,1683), 10 ng/mL Activin A (StemCell™, 78001.1), 20 ng/mL LIF (StemCell™, 78055).Approximately 6.6x10^5^ mouse embryonic fibroblasts were seeded onto each well of a 6-well plate coated with 0.1 % gelatin. Primed hESCs were dissociated with 30% TrypLE and seeded onto the MEF plates using mTESR-plus media supplemented with Y-27632. Two days after seeding hESCs, media was replaced with 5iLAF media to begin conversion of the primed hESCs to a naïve state. Cells were cultured in 5% O_2_, 5% CO_2_ hypoxia chambers and media was refreshed every two days. Naïve hESCs were then transdifferentiated by passaging cells into hTSCs media and sorted for ITGA2^+^ and EPCAM^+^ cell populations.

### Culture of first-trimester placental cytotrophoblasts and inhibitor treatment

First trimester (6-8 weeks) placental villi were collected after approval from the University of Kansas Medical Center Institutional Review Board. Cytotrophoblasts were isolated following established protocols from Okae et al. Placental villi were washed with PBS, followed by two consecutive digestions with 1x Hanks’ Balanced Salt Solution (Sigma-Aldrich^TM^, H4641). 0.125% trypsin (Gibco^TM^, 15090-046) and 0.125 mg/mL DNase I (Sigma-Aldrich^TM^, DN25) at 37 °C. CTBs were then isolated through Fluorescence Activated Cell Sorting of CD49f PE conjugated antibody (Miltenyi Biotec, 130-119-767). Isolated CTBs were cultured on collagen IV coated plates and media was refreshed every two days. CTBs were cultured in DMEM supplemented with 0.1 mM 2-mercaptoethanol, 0.2% fetal bovine serum, 0.5% penicillin-streptomycin, 0.3% bovine serum albumin, 1% ITS-X, 1.5 μg/mL L-ascorbic acid, 50 ng/mL epidermal growth factor, 2 μM CHIR99201, 0.5 μM A83-01, 1 μM SB431542. 0.8 mM Valproic acid and 5 μM Y-27632.

FOR DNMT1 inhibition analysis in CTBs, 1x10^5^ CTBs were cultured and treated with either DMSO or GSK-3484862 (Med Chem Express, HY-135146) for five to nine days. RNA from CTBs were isolated for RNA sequencing analysis. Libraries were generated using the NEBNext Ultra II Directional RNA library Prep Kit for Illumina using manufacturer’s instructions.

### Immunofluorescence staining of hTSC and hESCs

hTSCs and hESCs were cultured on 24-well plates with a coverslip that were either coated with collagen IV or Matrigel, respectively. Cells were then fixed with 4% paraformaldehyde and washed with PBS for 5 minutes twice. Fixed cells were then blocked and permeabilized using 5% donkey serum with 0.1% Triton-X for 15 minutes at room temperature. Cells were then washed with PBS + 0.1% Tween-20 (PBS-T) for 5 minutes twice. Primary antibody was added with 5% donkey serum and 0.1% triton-X to each slide and incubated overnight at 4 °C. Cell were then washed with PBS-T 5 minutes twice and secondary antibody (Alexa Fluor 488/594/647, Invitrogen) and DAPI was then added to the cells for 1 hour at room temperature. Cells were then washed twice with PBS-T and coverslips were then mounted on frosted microscope slides using ProLong™ Diamond antifade mounting media. Cells were then imaged on the Zeiss LSM710 confocal microscopes and analyzed on FIJI software.

The following antibodies were used for this study:

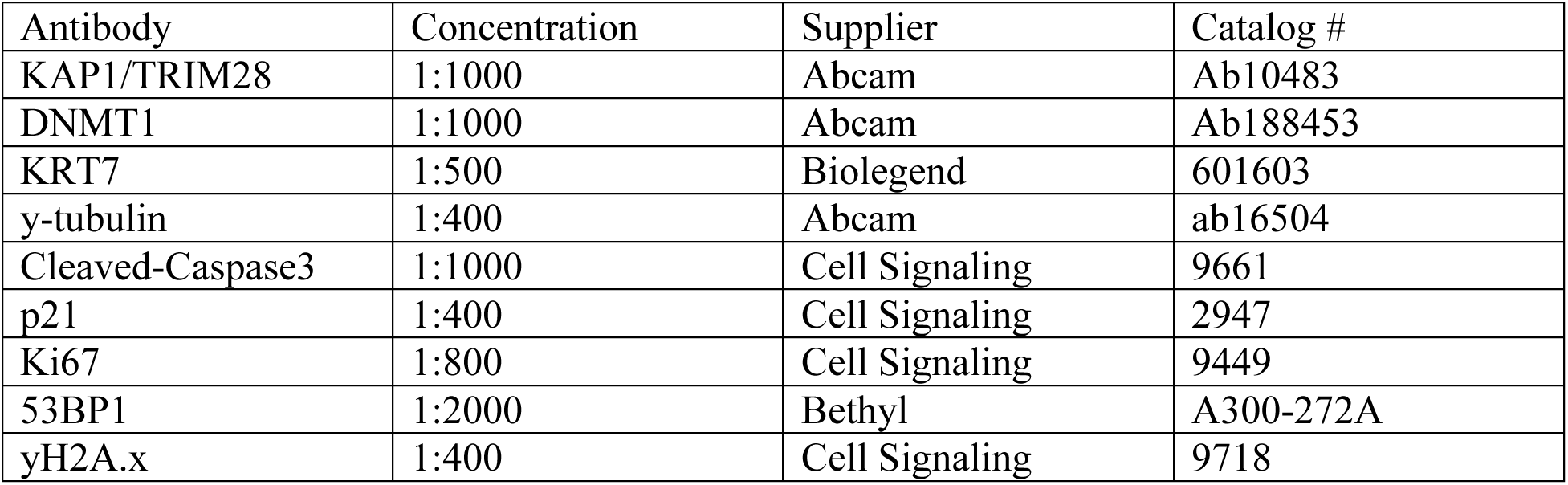

### Chromosomal Counting of DNMT1^Exon33/Exon33^ stem cells

Mutant and non-target bulk population hTSCs were treated with KaryoMAX^TM^ Colcemid^TM^ (Gibco^TM^, 15212012) solution supplemented into culture media at a concentration of 0.05 ug/mL for 1 hour. Cells were then dissociated with 30% TrypLE for 10 minutes and inhibited with Soybean Trypsin Inhibitor. The cells were then spun down and resuspended in 1.5 mL of fresh media. 10 mL of warmed 0.075 M potassium chloride (KCl) solution was added drop by drop while flicking tube to resuspend the cells. The cells were mixed by inverting the tube and placed into a 37 °C incubator for 10-15 minutes. Two to three drops of cold Carnoy’s fixative (3:1 methanol:glacial acetic acid) was then added to the cell suspension. Cells were then spun down for 5 minutes at 300 xg and the supernatant was aspirated leaving behind 1 mL to flick and resuspend the cells. 10 mL of cold Carnoy’s fixative was added drop by drop while gently vortexing the cells and then the cells were centrifuged for 5 minutes at 300 xg. This was repeated 3 times and samples were stored at 4 °C until all timepoints were collected. Microscope slides were then washed in cold Carnoy’s fixative. The cell suspension was resuspended in 500 uL of cold Carnoy’s fixative and 80 uL of cells were dropped from a height of 55 cm while holding the wet slide at an angle of 30° to allow the cells to explode and spread across the slide. Three to four drops of cells were added to each slide for a total of 4 slides per sample. A few drops of cold fixative were then added to each slide and then were then air dried for 10 minutes. Slides were then stained for DAPI and Human centromeric DNA FISH probes from PNA Bio (pan-centromere probe: AAACTAGACAGAAGCATT, catalog: F3006) by heating the slides at 85 °C for 5 minutes. Mix 0.2 uL (500 nM final concentration) of PNA probe and 100X DAPI with 20 uL hybridization buffer (20 mM Tris, pH 7.4, 60% formamide, 0.5% blocking reagent (Roche, 11096176001)). We then heated the hybridization buffer containing the probe 85 °C for 5 minutes and then added it to the slide and kept the slides heated at 85 °C for an additional 10 minutes. The slides were then incubated at room temperature for 2 hours in the dark. Next, we washed the slides in wash buffer (2x SSC + 0.1% Tween-20) at 60 °C for 10 minutes and once at room temperature. We then washed the slides with 2X SSC for 2 minutes, 1X SSC for 2 minutes, and water for 2 minutes. Coverslips with mounted with ProLong Diamond Mounting media and slides were imaged at 100x using oil immersion on the LSM710 Zeiss Microscope.

### Western Blot

Protein from either hTSC or hESCs were extracted using ice cold RIPA buffer supplemented with fresh protease inhibitors and Halt’s protease inhibitor. Cell pellets were then sheared using the Bioruptor Pico on the following mode, “Easy shear” for 5 minutes. Protein lysates were then centrifuged for 15 minutes, and the lysate was transferred to a new tube to remove cell debris. Protein lysates concentrations were then measured using bradford assay and 5 -50 ug of protein were then run on a SDS-PAGE gel at 100 volts for 1.5 hours. The resolved proteins were then transferred to a Immunoblot® FL PVDF membrane (Bio-RAD, 1620264) for 2 hours at 300 mA. The membranes were then blocked using LI-COR Intercept blocking buffer for one hour at room temperature. Primary antibody was then added overnight in 1x Intercept blocking buffer + 0.15% Tween-20 at 4^°^C. Membranes were then washed for 5 minutes twice using 20 PBS-T and then incubated with a secondary ?luorescent antibody (IRDye® 680RD Donkey anti-mouse, 1:20000 & IRDye® 800CW Donkey anti-rabbit, 1:20000) for 1 hour at room temperature. The membrane was then washed for 5 minutes twice in PBS-T and then kept in PBS before imaging on the LI-COR Odyssey imaging system.

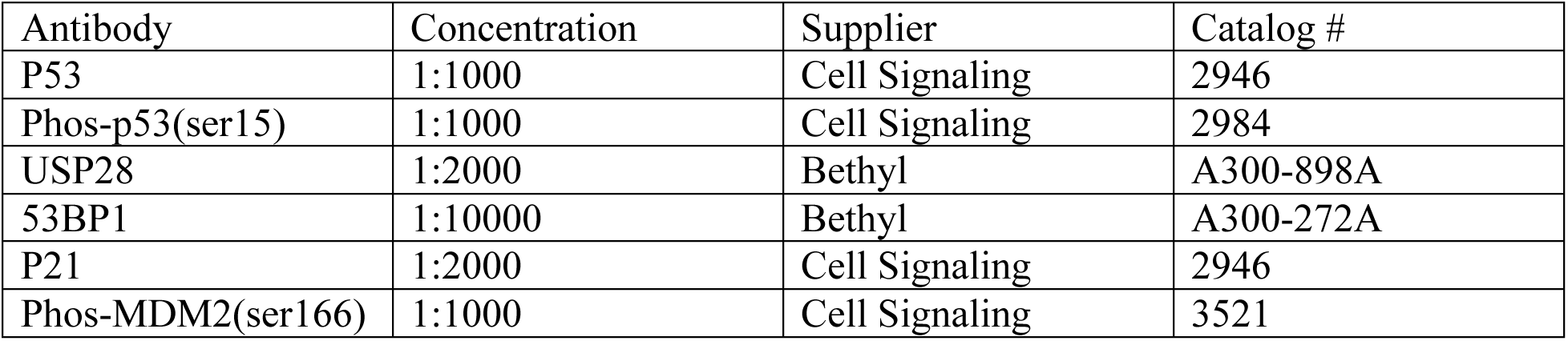

| Antibody | Concentration | Supplier | Catalog # |
| --- | --- | --- | --- |
| P53 | 1:1000 | Cell Signaling | 2946 |
| Phos-p53(ser15) | 1:1000 | Cell Signaling | 2984 |
| USP28 | 1:2000 | Bethyl | A300-898A |
| 53BP1 | 1:10000 | Bethyl | A300-272A |
| P21 | 1:2000 | Cell Signaling | 2946 |
| Phos-MDM2(ser166) | 1:1000 | Cell Signaling | 3521 |

### Flow Cytometry for cell cycle analysis

To measure cell cycle analysis of both *DNMT1^+/+^* and *DNMT1^-/-^* hTSCs, we harvested hTSCs at Day 4, 7 and 10 using TrypLE and Trypsin inhibitor. Cells were then resuspended in 300 ul of PBS and 100% ice-cold ethanol was then added to the cell drop-wise to slowly to fix the cells in a final concentration of 70% ethanol. The cells were then placed at -20 °C until all the samples were collected. Cells were spun down at 300 xg for 10 minutes and the 70% ethanol was aspirated. Cells were then washed with PBS once and resuspended in PBS with proprium iodide. All samples were measured on the FACS Aria flow cytometry machine using PE filter and analysis was done on FlowJo version 10.

### RNA isolation, cDNA generation and qPCR analysis

To isolate mRNA, 150 μL of RNAzol® RT (Molecular Research Center, RN 190) was added to each cell pellet and mixed. 60 μL of RNase-free water was then added to each sample. Samples were then vortexed and incubated for 10 minutes at room temperature. Samples were then centrifuged at 15000 xg for 15 minutes at 4 °C and the supernatant was collected. mRNA was then precipitated with 80 μL of 75 % ethanol. Samples were then vortexed, incubated for 10 minutes at room temperature and centrifuged at 15000 xg for 15 minutes at 4 °C. The supernatant was aspirated, and the pellet was washed with 500 μL of 75% ethanol twice. Finally, the supernatant was then aspirated and samples were left to air dry for 10 minutes and resuspended in 30 μL of RNase-free water.

500 ng of isolated RNA was used to generate complementary DNA using the SensiFAST cDNA synthesis Kit (Bioline, BIO65053) using manufacturer’s instructions. cDNA from samples were diluted 10 ng/μL and used to set up the qPCR reaction with PowerUp SYBR using manufacturer’s instructions. The qPCR reaction mix was then placed into the QuantStudio5 qPCR thermocycler machine.

The following primers were used for qPCR analysis:

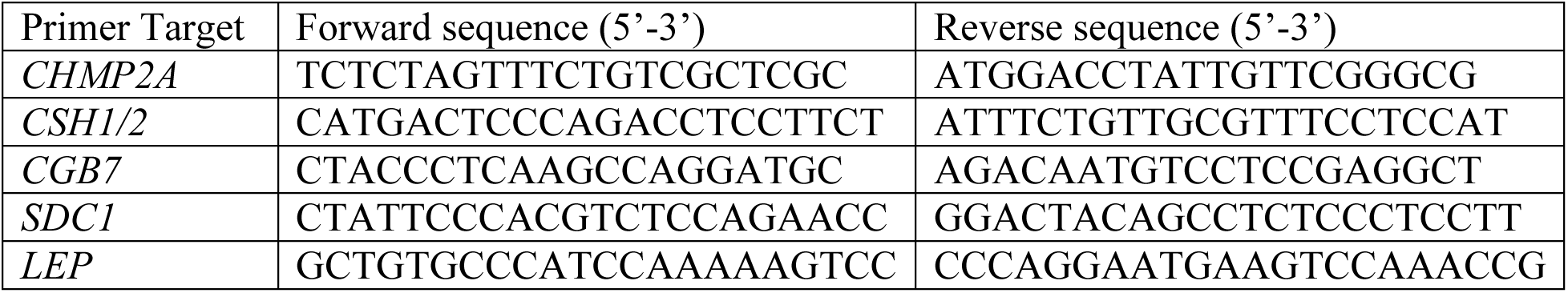

### Whole Genome Bisulfite Sequencing (WGBS) library generation

*DNMT1^-/-^* hTSCs were harvested using TrypLE and Trypsin inhibitor and then cell pellet were flash frozen with liquid nitrogen. Genomic DNA from cell pellets were isolated using the QiaAMP DNA mini kit (Qiagen, 51304) and quantified using the Qubit fluorometer. 500 ng of isolated genomic DNA with 1.25 ng of lambda DNA spike-in was sheared using the Covaris M220 ultrasound sonicator to an average sheared size of 350 bp. 200 ng of sheared DNA was then bisulfite converted using the EZ DNA Methylation-Gold Kit (Zymo Research, D5005) using manufacturer’s instructions. 100 ng of bisulfite converted DNA was then used to generate WGBS libraries using the Swift Accel-NGS® Methyl-Seq DNA Library Kit (Swift Biosciences, 30024) according to manufacturer’s instructions. The final concentrations of the WGBS libraries were determined using the Qubit 1x DNA high-sensitivity assay kit.

### Whole Genome Bisulfite Sequencing analysis

Measurement of DNA methylation was done on *DNMT1^-/-^* hTSCs in this project and used to compare DNA methylation between hTSCs, CTBs, Primed and Naïve hESCs from public datasets. WGBS reads were trimmed using Trim Galore! (v0.6.6) (https://github.com/FelixKrueger/TrimGalore) with the following parameters (-q 20, - clip_R1 5, -clip_R2 10) to remove the first 5 bases of read 1 and 10 bases of read 2. The trimmed fastq files were then aligned to the hg38 human reference genome using Bismark (v0.23.0)^27^ using default parameters. The resulting aligned BAMs were then deduplicated and filtered for incomplete bisulfite conversion using Bismark (v0.23.0). Methylation over cytosines were calculated using the methylation call function in Bismark. Metaplot and heatmap analysis of CpG methylation over MER11A regions were measured by calculating the average CpG methylation over the start site and 3 Kb flanking regions. In addition, CpG methylation over DNA methylation sensitive genes in hTSC and hESCs were filtered from significantly upregulated genes in *DNMT1^-/-^*cells and average CpG methylation was measure from -700 bp to 300 bp of the transcription start site of each gene.

### RNA isolation and RNA sequencing

Mutant and non-target bulk population hTSCs and hESCs were collected for total RNA isolation using the Qiagen RNAeasy Mini Kit using manufacturer’s instructions. Total RNA was then quantified using the Qubit and RNA integrity was validated though gel electrophoresis or the Bioanalyzer. RNA sequencing libraries were then generated using the NEB Poly(A) mRNA Magnetic Isolation Module and the NEBNext Ultra II Directional RNA library Prep Kit for Illumina using manufacturer’s instructions.

### RNA sequencing analysis

Both samples sequenced for this project and publicly downloaded fastq were first trimmed using trimmomatic (v0.34)^28^ to remove contaminated adapters and low-quality reads. The resulting trimmed fastqs were then aligned to the hg38 human reference genome using STAR (v2.7.8a)^29^ with default parameters. Picard (v2.9.0) (https://broadinstitute.github.io/picard/) was then used to sort and mark duplicate reads from aligned bam files and the resulting reads were then quantified using HTseq-count^30^ and the StringTie suite (v1.3.5)^31^. Read counts for each sample were then used to measure differential gene expression patterns using DESeq2^32^. ClusterProfiler^33^ was used for GOTERM analysis and GSEA (PMID: 16199517) for p53-response genes was used using default parameters.

The TEtranscript pipeline^34^ was used to assess difference in transposon class expression in *DNMT1^-/-^*hESCs and hTSCs and *TRIM28^-/-^* hESCs and hTSCs. Fastqs for each file were trimmed using Trim Galore! (v0.6.6) using default parameters. The trimmed fastqs were then aligned to the hg38 Human reference genome using STAR with parameters that support multiple alignments per read (–winAnchorMultimapNmax 200,-outFilterMultimapNmax 100). The aligned BAM files were then used to generate counts for each transposon class and DESeq2 was used to measured differential expression of each of these transposons.

### Chromatin Immunoprecipitation

hTSCs were cultured until close to confluency in a single well of a 6-well plate (approximately 1.2x10^6^ cells) and were crosslinked with 1% paraformaldehyde for 15 minutes. The fixed samples were then quenched with 1 mM glycine for 10 minutes. Cells were then lysed using lysis buffer solution and sonicated with the M220 ultrasonicator (Covaris) in a 1 mL AFA Fiber tube using the following conditions (cycles/burst = 200, duty factor = 20%, peak intensity = 75, time = 10 minutes and temperature = 7 °C). The resulting sheared DNA was pre-cleared using magnetic beads (Sera-Mag Protein A/G SpeedBeads, VWR 17152104010150) for 2 hour to remove non-specific binding by the antibody. The beads were then removed using a magnetic rack and the sonicated lysate was transferred to a new tube and 1 μL of TRIM28 antibody was added to the sample and incubated overnight. The following day, magnetic beads were added to the sample and incubated for 2 hours at 4 °C to allow the beads to bind to the TRIM28/DNA complex. Samples were then washed with buffer of increasing salt concentration (Wash Buffer A: 50 mM HEPES, 1% Triton X-100, 0.1% deoxycholate, 1 mM EDTA, 140 mM NaCl and Wash Buffer B: 50 mM HEPES, 0.1% SDS, 1% Triton X-100, 0.1% deoxycholate, 1 mM EDTA, 500 mM NaCl) and TE buffer). The TRIM28/DNA complex was then eluted from the beads using elution buffer (50 mM Tris–HCl, 1 mM EDTA, 1% SDS) incubated at 65 °C for 10 minutes and placed on a magnetic rack. The supernatant was then transferred to a new tube and de-crosslinked by incubating at 65 °C overnight. Residual RNA and protein was removed from the sample by adding 1.5 μL of 10 mg/mL RNaseA, incubated for 30 minutes at 37 °C, and 10 μL of 10 mg/mL Proteinase K, incubated for 2 hours at 56 °C, to each sample. DNA was then purified using the Qiagen MinElute PCR Purification Kit using manufacturer’s instructions.

### Chromatin Immunoprecipitation data analysis

Original and publicly available datasets for ChIP and CUT&RUN were used in this paper (Table S2). Downloaded fastq files were trimmed using trimmomatic (v0.6.6) using default parameters and the resulting trimmed fastq were then aligned to the hg38 Human reference genome using BWA (v0.7.17)^35^. To account for repetitive regions in the genome, the aligned BAMs were not filtered for unique reads. Peak calling for each respective ChIP and CUT&RUN experiment was done through MACS2^36,37^ using default parameters and inputs as a control and significant peaks were corrected for false discovery rate using the Benjamini-Hochberg correction. Bigwigs of aligned samples were made through DeepTools^38^ and were used to generate metaplots and heatmaps using the computeMatrix, plotHeatmap and plotProfile functions. Normalization of ChIP metaplots were calculated by (1) measuring the median signal of the ChIP signal throughout the genome, (2) then measuring the max height of the signal peaks across all genetic elements of interest (if look at specific promoters, we measure the max height across all promoters), (3) a scaling factor was then calculated by subtracting the height from (2) and the median signal from (1). (4) Specific ChIP signal was then normalized across regions of interest by the following equation: (signal height – median)/scaling factor. Region Associated DEGs (RAD) analysis^39^ algorithms were used to measure the relationship of ChIP peaks to proximal DEGs.

### ATAC data analysis

Publicly available datasets for ATAC-seq were used in this paper (Table S2). Downloaded fastq were trimmed using trimmomatic (v0.6.6) with default parameters. The resulting trimmed fastq files were then aligned with the hg38 Human reference genome using BWA (v0.7.17) and were not filtered for the unique reads to account for repetitive regions. Bigwigs of the aligned reads were made using DeepTools including any metaplot/heatmap analysis using the computeMatrix and plot/Heatmap/plotProfile functions.

## RESULTS

### Loss of TRIM28 and gain of DNA methylation at transposons in concert with developmental progression

To study the human placental and embryonic lineage, we used human TSCs (hTSCs) cells and hESCs respectively. hTSC are derived from first-trimester placental cytotrophoblasts (CTBs)^24^. hTSCs show analogous methylation patterns to CTBs, except that large regions hypomethylated in CTBs, called partially methylated domains, show even greater hypomethylation in hTSCs^5,24,40^. hESCs can be cultured in naïve and primed culture conditions, resulting in a transcriptional state and global 5mC level analogous to pre- and post-implantation epiblast respectively^41,42^. Naïve hESCs can be converted to hTSCs and primed hESCs and are the most primitive population of the three^20,43,44^.

To assess possible mechanisms of transposon silencing in human epiblast and trophoblast lineage, we conducted ChIP-seq for TRIM28 in hTSCs and mined TRIM28 ChIP-seq for naïve and primed hESCs^45^. We made use of published ATAC-seq^46,47^, and DNA methylation data from hTSCs^24^ and naïve and primed hESCs^42^, assigning enrichment for these features across subclasses of long terminal repeats (LTRs), the regulatory elements which flank endogenous retrovirus transposons^48^ (Tables S1 – S3). We also mined DNA methylation data from human cytotrophoblasts (CTBs)^10^, whose methylation pattern correlated fairly well with hTSCs over LTRs (r^2^ = 0.696) (Figure S1A-E).

Consistent with TRIM28 being the predominant transposon-silencing mechanism in early embryonic development, far more TRIM28 sites were detected in naïve hESCs than in primed hESCs and hTSCs (Figure 1A,B). This is not a result of differences in ChIP-seq quality or cut-off effects, as many clearly defined TRIM28 sites in naïve hESCs are absent in primed hESCs and hTSCs (Figure 1B-E, S1F). The numerically most abundant TRIM28 sites in naïve hESCs correspond to THE1B and LTR12C, transposon classes that function as regulatory elements during zygotic gene activation (ZGA) during the 8-cell stage of development and are silenced by the epiblast stage^49,50^ (Figure 1C). TRIM28 is lost from these elements in the more developmentally advanced primed hESC and hTSC (Figure 1B-G). THE1B elements appear to be generally transcriptionally inert in the cell types studied, but the CpG-rich LTR12C elements are specifically targeted for DNA methylation in the more developmentally advanced cells (Figure 1H, S1B-E) and show concomitantly lower chromatin openness (Figure 1I). A similar process of replacement of TRIM28 with 5mC is observed at other transposon classes, notably SVA elements (Figure S1G-I). Not all transposon classes lose TRIM28 though. Most strikingly, MER11A and B sites retain a high degree of TRIM28 binding in primed hESCs and hTSCs (Figure 1C-E,J,K).

**Figure 1.**
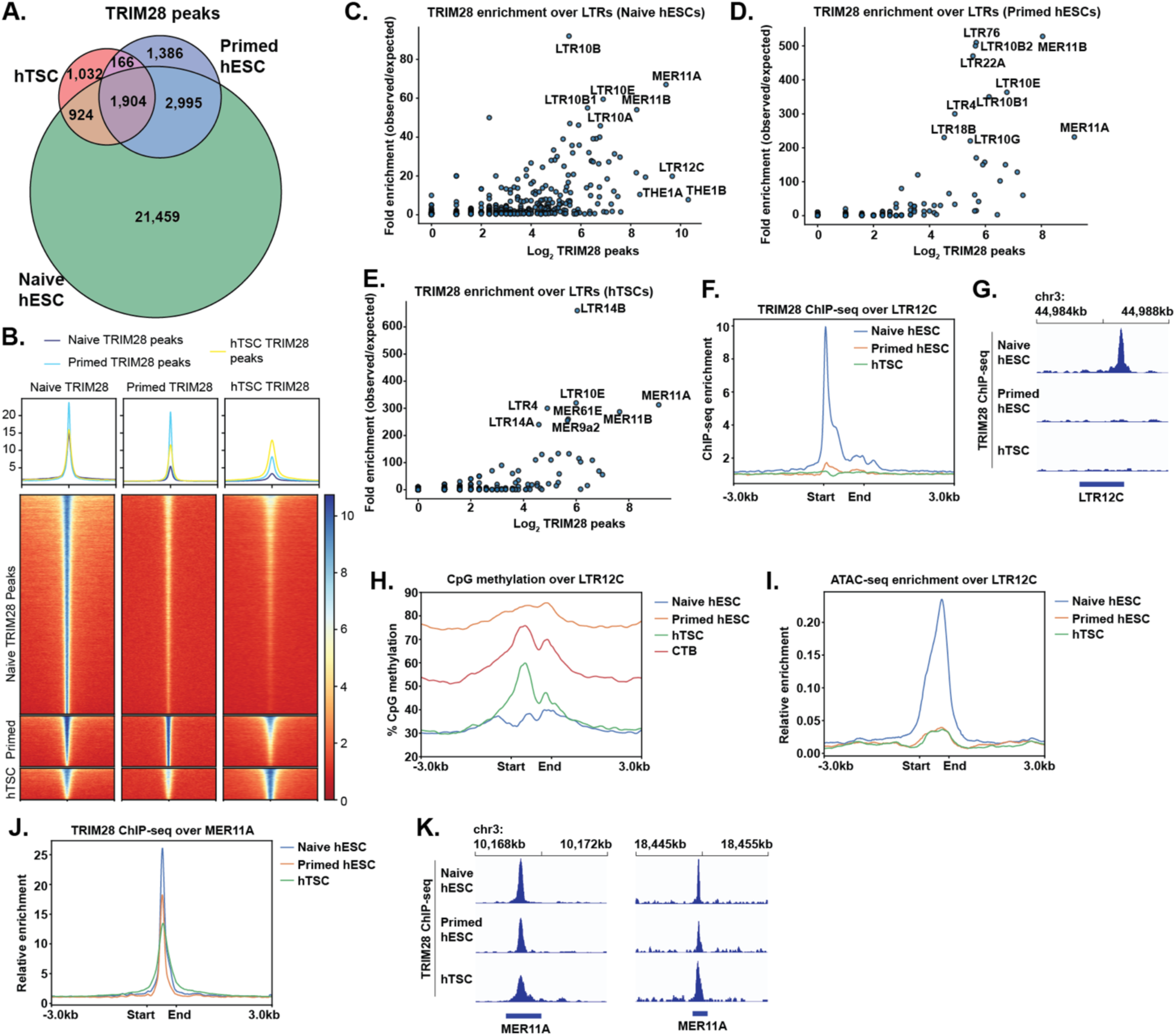
Distribution of TRIM28 and 5mC over transposons in embryonic and trophoblast stem cells. (A) Venn diagram of TRIM28 peaks in naïve hESCs, primed hESCs and hTSCs. (B) Heatmap of TRIM28 enrichment over peaks present in naïve hESCs, primed hESCs, and hTSCs. (C – E) Number of TRIM28 peaks overlapping LTR classes, and fold enrichment TRIM28 peaks over the LTR class in question, in naïve hESCs (C), primed hESCs (D), and hTSCs (E). (F) Metaplot of TRIM28 enrichment over all LTR12C elements in cell types indicated. (G) TRIM28 enrichment over a representative LTR12C element. (H) Metaplot of %CpG methylation over all LTR12C elements in cell types indicated. (I) Metaplot of ATAC-seq enrichment over all LTR12C elements in cell types indicated (J) Metaplot of TRIM28 enrichment over all MER11A elements in cell types indicated. (K) TRIM28 enrichment over two representative TRIM28 elements.

Despite limited TRIM28 distribution and low global DNA methylation (Figure S1A), hTSC did not show generally elevated levels of transposon expression relative to hESCs (Figure S1J-M), ruling out the possibility that hTSCs simply have less control of transposons.

### TRIM28 regulates gene expression by suppressing adjacent enhancers

To determine TRIM28’s regulatory role in trophoblast, we knocked out *TRIM28* in primed hESCs and hTSCs, using two sgRNA to excise an early exon and induce a frameshift (Figure S2A). *TRIM28^-/-^* primed hESCs could survive and self-renew as previously reported^51^ (Figure S2B), but we were unable to recover *TRIM28^-/-^* hTSCs clonal lines. Four days after nucleofection of Cas9 protein loaded with sgRNA targeting *TRIM28*, almost all hTSC showed loss of TRIM28 protein, but the residual TRIM28-expressing cells eventually took over the culture (Figure 2A-C), further indicating that *TRIM28^-/-^* hTSCs cannot survive long term in culture. We thus conducted RNA-sequencing of bulk TRIM28 KO and nucleofected control populations at four and seven days after nucleofection to identify TRIM28 regulatory targets (Figure 2A, Table S4).

**Figure 2.**
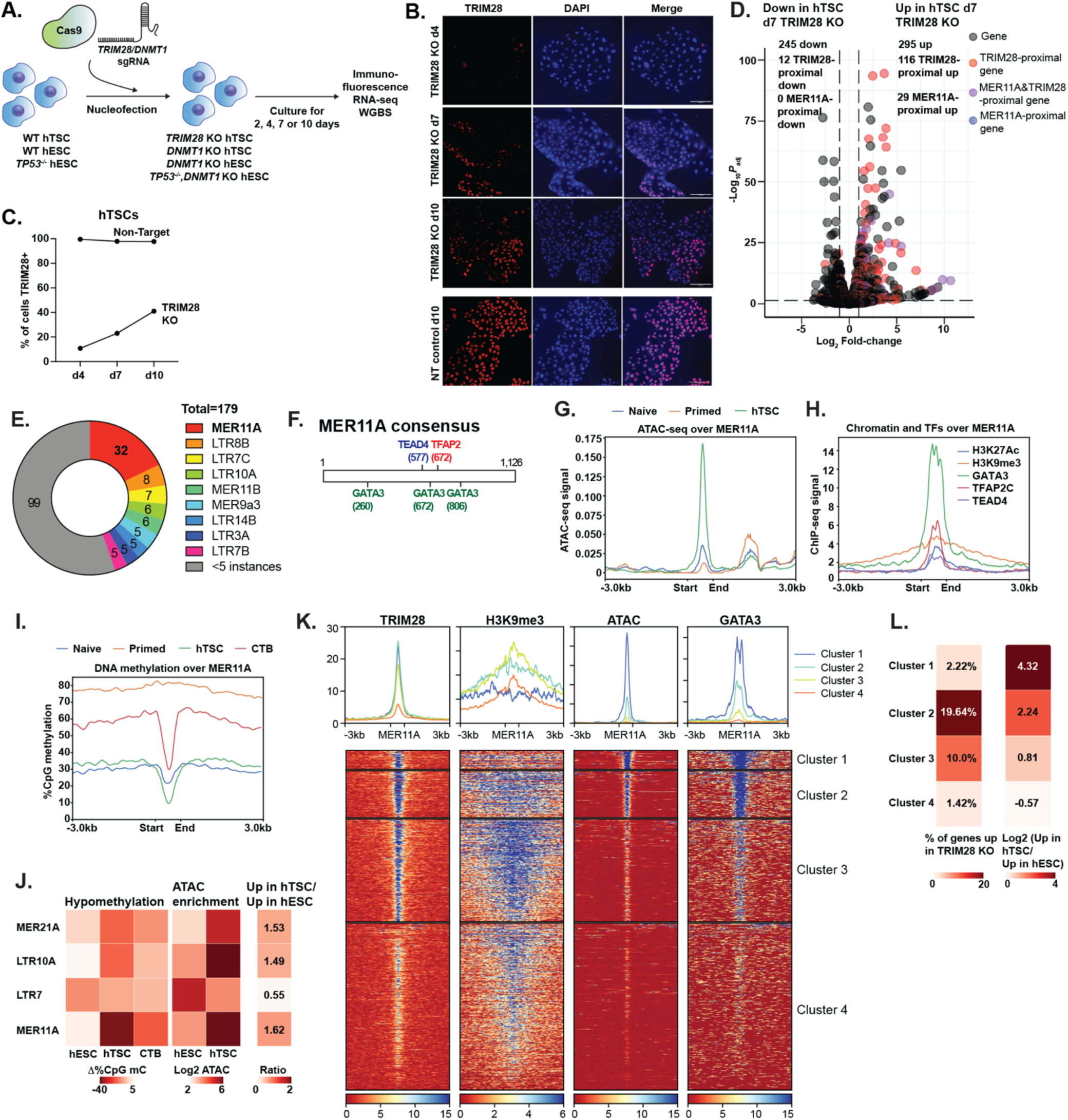
TRIM28 regulates adjacent genes by suppression of LTR transposons. (A) Schematic of experiments in which DNMT1 or TRIM28 is ablated via CRISPR/Cas9 nucleofection and phenotypic changes are subsequently observed in the bulk population of cells. (B) Immunofluorescent staining for TRIM28 in a bulk population of hTSCs 4,7 or 10 days after nucleofection with CRISPR/Cas9 and sgRNA targeting TRIM28 or a non-targeting control sgRNA. (C) Percentage of hTSC positive for TRIM28 after nucleofection with CRISPR/Cas9 and sgRNA targeting TRIM28 (TRIM28 KO) or non-targeting sgRNA (Non-target) (D) Volcano plot comparing bulk population of TRIM28 KO hTSC to control, with TRIM28 (red), MER11A- (blue), TRIM28 & MER11A-proximal genes (purple) within 50 kb highlighted. 3 upregulated and one downregulated gene were off-axis and not plotted. (E) Pie-chart of TRIM28-bound LTRs within 50 kb of a gene upregulated in TRIM28 KO hTSCs. (F) Schematic showing location of TEAD4, TFAP2C, and GATA3 motifs over a MER11A consensus element. (G) Metaplot of ATAC-seq signal over MER11A elements in cell types indicated. (H) Metaplot of ChIP-seq signal from chromatin mark or transcription factor indicated over MER11A elements in hTSCs. (I) Metaplot of CpG methylation over MER11A elements in cell types indicated. (J) Heatmap showing extent of localized hypomethylation, enrichment for ATAC-seq signal, and likelihood of being upregulated in hTSC vs. hESC for the LTR classes indicated. (K) Heatmap showing k-means clustering of MER11A elements according to TRIM28, H3K9me3, ATAC-seq and GATA3 binding. A fifth cluster of 4 MER11A elements showing extremely high signal for all features is not plotted. (L) Percentage of genes upregulated in TRIM28 KO hTSC vs. control hTSC, or hTSC vs. hESC-specific genes, for each cluster in (K).

Neither TRIM28 KO hTSCs nor published data from *TRIM28^-/-^*hESCs showed a global increase in the fraction of RNA-seq reads derived from transposons (Figure S2C,D)^51^. We found however that genes upregulated in the TRIM28 KO were disproportionately likely to be proximal (<50kb distance) to TRIM28 sites (Figure 2D). The TRIM28 sites near upregulated genes were associated with a variety of LTR elements, with MER11A by far the most frequent (Figure 2E, Table S5). Upregulated transcripts did not originate from MER11A (Figure S2E, S2F), indicating an enhancer rather than promoter function for these LTRs upon TRIM28 loss.

Genes upregulated in TRIM28 KO naïve and primed hESCs also showed proximity to transposon-associated TRIM28 sites, (Figure S2G, S2H), suggesting a similar mechanism of TRIM28-mediated suppression of LTR enhancer activity.

### Putative enhancer function for MER11A elements in WT hTSCs

The MER11A consensus sequence contains motifs for the placental transcription factors GATA3, TEAD4, and TFAP2C (Figure 2F). On average, MER11A sites showed elevated openness in hTSCs (Figure 2G), enrichment for both heterochromatic and euchromatic marks (Figure 2H), and pronounced hypomethylation in hTSCs and CTBs, which is a hallmark of enhancers^52^ (Figure 2I). This raised the interesting possibility that even with TRIM28 intact, MER11A elements have been co-opted as enhancers in the trophoblast lineage, akin to other transposon classes such as MER21A, MER41A/B, MER39B and LTR10A^12,13^. Genes proximal to MER11A elements were indeed preferentially upregulated in hTSCs relative to hESCs to an extent similar to the classes above, while genes proximal to the hESC-specific LTR7 class^45^ showed the opposite trend (Figure 2J).

We performed cluster analysis of chromatin features of MER11A elements and identified four distinct clusters (Figure 2K). Cluster 1 sites were primarily euchromatic, with strong ATAC-seq and GATA3 enrichment. Cluster 3 showed strong enrichment for H3K9me3, a heterochromatic mark deposited by the TRIM28-associated histone methyltransferase SETDB1^8^. Cluster 2 showed simultaneous enrichment for eu- and heterochromatin features. Cluster 4 elements showed weak enrichment for all tested marks and greater divergence from the MER11A consensus sequence. Appropriately, genes proximal to Cluster 1 and 2 sites were more likely to be upregulated in hTSC relative to hESCs, while genes proximal to Cluster 2 and 3 sites showed upregulation upon TRIM28 KO (Figure 2L). Six KRAB-ZFN proteins, capable of TRIM28 complex recruitment, are enriched over MER11A sites based on ChIP-seq data in HEK293T cells^53^. Five of these KRAB-ZFNs showed similar enrichment over Clusters 1 – 3, but ZFN468 showed lower enrichment over Cluster 1, potentially accounting for the weaker silencing of these loci (Figure S2I).

Interestingly, sequences from the MER11A element proximal to the *LEP* gene have already been demonstrated to show reporter activity selectively in choriocarcinoma (placental cancer) cells^54,55^, and high levels of placental leptin occur in old-world but not new-world primates^56^, corresponding to the presence of the MER11A element at the locus (Figure S2J,K). CRISPRi targeting of this element resulted in suppressed *LEP* expression in hTSCs and differentiated syncytiotrophoblast (STBs), without any effect on expression of STB differentiation markers (Figure S2L-P). In total, some MER11A elements serve as putative enhancers that regulate transcription in normal hTSCs, while others are suppressed by TRIM28.

### DNA methylation is essential for survival and transposon repression in hESCs and hTSCs

To determine the regulatory role of DNA methylation, we ablated the maintenance methyltransferase *DNMT1* in hESCs and hTSCs, using two sgRNA to ablate a critical exon in the catalytic domain (Figure S3A). Loss of *DNMT1* in hTSC or primed hESC was lethal, with high efficiency deletion followed by emergence of residual DNMT1-expressing cells (Figure 3A-C, S3B-C). After DNMT1 deletion, global DNA methylation in hTSCs dropped and then rebounded as residual DNMT1-expressing cells took over the bulk population (Figure 3D).

**Figure 3.**
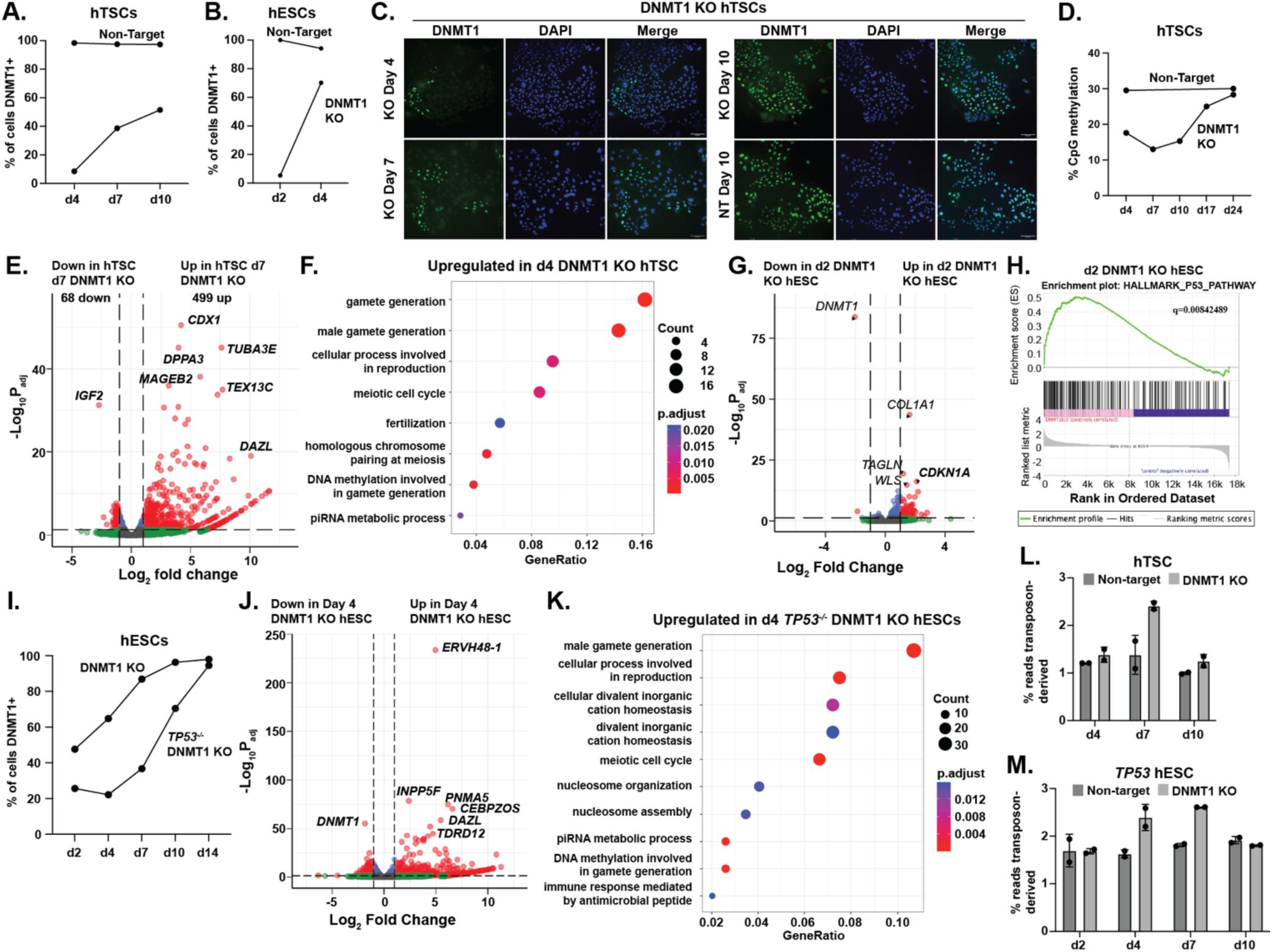
DNA methylation is essential for survival and transposon repression of hTSCs and hESCs. (A-B) Percentage of hTSC (A) or hESC (B) positive for DNMT1 after nucleofection with CRISPR/Cas9 and sgRNA targeting DNMT1 (DNMT1 KO) or non-targeting sgRNA (Non-target) (C) Immunofluorescent staining for DNMT1 in a bulk population of hTSCs 4 to 10 days after nucleofection with CRISPR/Cas9 and sgRNA targeting DNMT1. hTSCs that underwent nucleofection with a non-targeting sgRNA are shown as a control. (D) Global CpG methylation in bulk population of hTSC targeted with CRISPR/Cas9 and sgRNA targeting DNMT1 (DNMT1 KO) or non-targeting sgRNA (Non-target) (E) Volcano plot comparing expression of hTSCs 7 days after targeting with DNMT1 sgRNA compared with cells nucleofected with non-targeting sgRNA. (F) Gene ontology analysis of genes upregulated in DNMT1 KO bulk population hTSCs four days after nucleofection relative to cells undergoing control nucleofection. (G) Volcano plot comparing expression of hESCs 2 days after targeting with DNMT1 sgRNA compared with cells nucleofected with non-targeting sgRNA. (H) GSEA analysis comparing expression of P53 pathway genes in day 2 DNMT1 KO hESCs and controls. (I) Percentage of WT and *TP53^-/-^* hESCs positive for DNMT1 between 2 and 14 days after nucleofection with DNMT1 sgRNA. (J) Volcano plot comparing expression of hESCs 4 days after targeting with DNMT1 sgRNA compared with cells nucleofected with non-targeting sgRNA. (K) Gene ontology analysis of genes upregulated in *TP53^-/-^* DNMT1 KO hESCs four days after nucleofection relative to *TP53^-/-^*hESCs undergoing control nucleofection (n=2 replicates). (L,M) Percentage of transposon-derived reads in hTSC (L) and *TP53^-/-^* hESCs (M) after nucleofection with sgRNA targeting DNMT1 or non-targeting sgRNA (n=2 replicates per genotype and timepoint).

We conducted RNA-seq in the bulk *DNMT1* KO population at timepoints following genetic deletion but before most knockout cells were lost. Consistent with DNA methylation’s established role in repression of germline genes, *DNMT1* KO hTSCs showed strong upregulation of germline genes (Figure 3E,F). *DNMT1* KO hESCs disappeared from culture extremely rapidly. Limited genic dysregulation was observed at day two and almost all KO cells were lost by day four (Figure 3B,G, S3C). One of the most strongly upregulated genes in day two DNMT1 KO hESCs was the p53 target gene *CDKN1A,* which encodes the tumour suppressor p21. The day two DNMT1 KO hESCs showed a general increase in expression of p53 pathway genes (Figure 3G,H), a phenomenon not observed in DNMT1 KO hTSCs (Figure S3D). Because rapid-p53 mediated loss made it difficult to determine DNMT1 regulatory targets, we generated *TP53^-/-^* hESCs (Figure S3E). These survived loss of *DNMT1* far longer and showed upregulation of germline genes (Figure 3I-K, S3F).

Loss of *DNMT1* results in widespread transposon upregulation and an increase in the total percentage of reads emanating from transposons in both hTSCs and hESCs (Figure 3L,M, S3G,H). DNA methylation appears to be the predominant transposon silencing mechanism in both hTSCs and primed hESCs cell types. Consistent with the replacement of TRIM28 with 5mC upon developmental progression described above, LTR12C, SVA, and LINE elements show a shift from TRIM28-mediated to 5mC-mediated repression in more developmentally advanced cells (Figure S3I,J).

### General and lineage-specific regulation in hESC and hTSC by DNA methylation

To identify the shared and lineage-specific genic targets of DNA methylation, we compared genes upregulated in d7 DNMT1 KO hTSCs and d7 DNMT1 KO *TP53^-/-^* hESCs relative to controls. A total of 223 genes were upregulated in both sets (Figure 4A, Table S6). As predicted, this gene set had far higher levels of promoter methylation than the set of all genes in trophoblast and epiblast-derived tissues and was strongly enriched for germline genes (Figure 4B, C). DNMT1 KO also caused upregulation of a set of genes specific to hTSC or hESC (Figure 4A). These genes also showed higher promoter methylation than the genome as a whole but included a significant subset of unmethylated genes which may be indirect targets (Figure 4D, E). To identify genuine examples of genes selectively regulated by DNA methylation in embryonic or placental lineage, we selected for genes which were 1) selectively upregulated by DNMT1 KO in one lineage 2) methylated in stem cells in that lineage 3) methylated *in vivo* in that lineage and 4) selectively methylated in the lineage in question (Figure S4A,B). This yielded a list of 22 putative methylation targets in hTSC and 120 targets in hESC, including 21 and 65 protein-coding genes respectively, including a handful of developmentally significant proteins (Figure 4F, S4C, Table S6).

**Figure 4.**
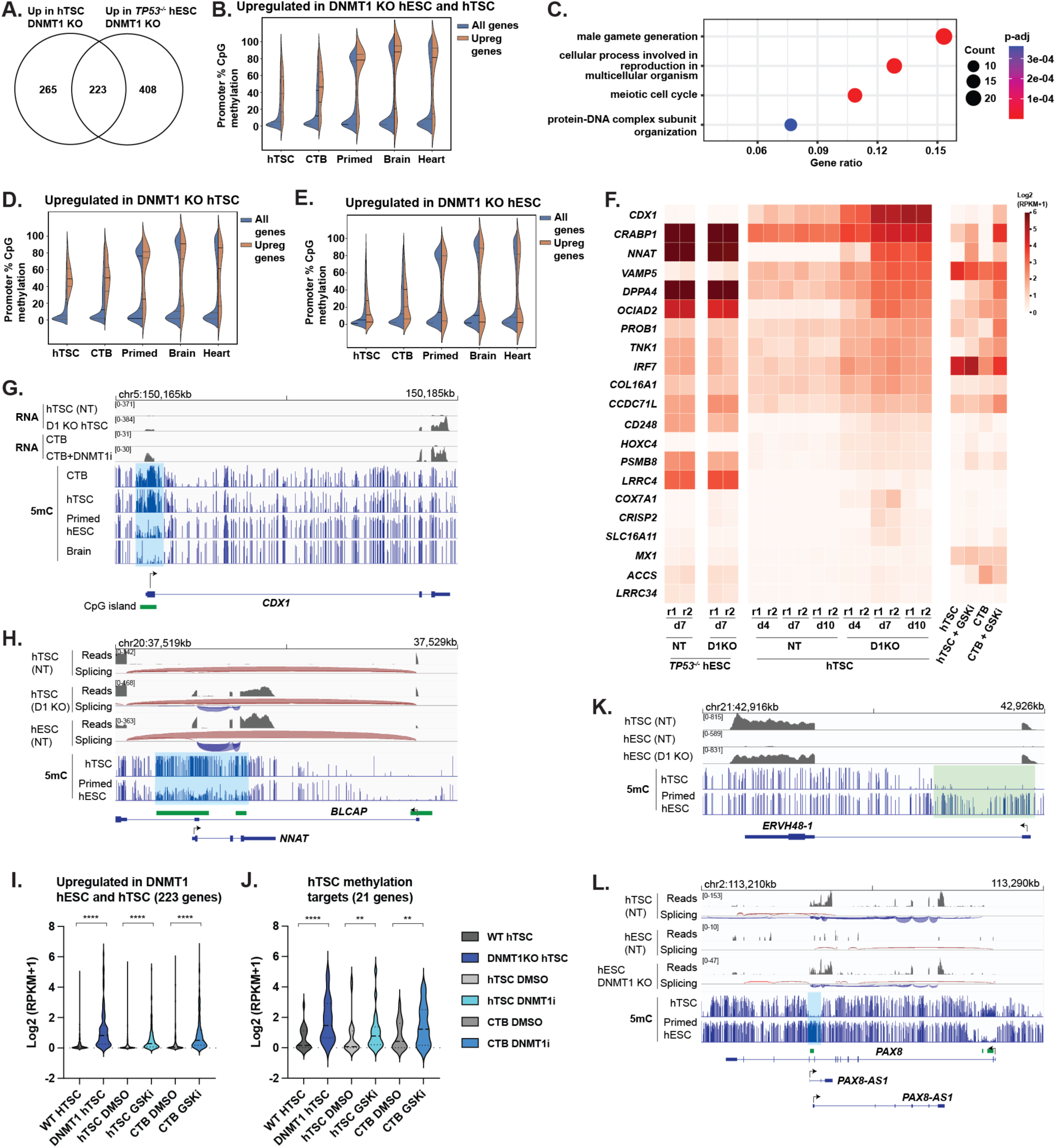
Regulation of developmentally important genes by DNA methylation in trophoblast and epiblast lineage. (A) Venn diagram of genes upregulated in DNMT1 KO hESCs and hTSCs. Genes in Venn diagram show fold-change >2, p_adj_<0.05 in at least one cell type, intersection indicates fold-change >2 in both cell types. (B) DNA methylation over the promoter of overlapping set from (A) in cell types indicated. (C) Gene ontology analysis of overlapping set in (A). (D, E) DNA methylation over the promoter of genes upregulated selectively in hTSCs (D) or hESCs (E) in (A). (F) Heatmap showing expression of genes identified as repressed by 5mC in hTSCs. (G,H) Methylation and transcription over the *CDX1* (G), *BLCAP/NNAT* (H). In H, splicing is also indicated: plus strand blue, minus strand red. (I,J) Expression of genes upregulated upon DNMT1 loss in both hESC and hTSC (I), or genes identified specifically as methylation targets in hTSC (J) in DNMT1 KO and DNMTi treated hTSCs and CTBs ****: p< 0.0001, **: p< 0.01 (K,L) Methylation and transcription over the *ERVH48-1* (K), *PAX8/PAX8-AS* (L) loci. In L, splicing is also indicated: plus strand blue, minus strand red.

Reflecting the differing methylation patterns in trophoblast and epiblast, genes regulated by hTSC-specific methylation are far more likely to contain CpG-island promoters (19/21 vs. 13/65 in hESCs). The most statistically upregulated gene in DNMT1 KO hTSCs was the transcription factor *CDX1* (Figure 3E, Figure 4G), which regulates anterior/posterior embryonic patterning and promotes intestinal cell fate^57,58^ and which is not normally expressed in any trophoblastic tissue. The pluripotency-associated transcription factor *DPPA4*^59^, which promotes H3K4 methylation at bivalent genes and thus antagonizes CpG island methylation in ESCs^60,61^, is itself strongly repressed by DNA methylation in trophoblast. Additional upregulated genes include *CRABP1*, which controls differentiation pathways by sequestering retinoic acid^62^, and *NNAT* (Neuronatin) a regulator of intracellular calcium^63^ and the only known protein-coding gene that is imprinted in somatic cells but fully methylated in placenta (Figure 4H)^64^. Both of these genes are highly expressed in human post-implantation epiblast, but not trophoblast (Figure S4D)^65^. To further validate of these targets, we treated primary CTBs with the DNMT1 inhibitor GSK-3484862^66^. We observed general upregulation of common methylation targets (Figure 4I) as well as specific upregulation of genes repressed by DNMT1 in hTSC (Figure 4F,G,J).

The MHC class I gene *HLA-A*, which must be silenced in placenta to prevent maternal immune rejection^67^, fell just below the threshold of CTB methylation to be included as a placental methylation target in our algorithm, but it shows strong evidence of suppression by 5mC. DNTM1 KO hTSCs and DNMT1i-treated hTSCs and CTB showed strong upregulation of HLA-A (Figure S4E). Analysis of published data further shows that blastocyst-derived hTSC (as opposed to placenta-derived) and at least some hTSC derived from hESC show leaky expression and aberrant hypomethylation of the HLA-A promoter (Figure S4F-H). Not only is it a gene whose silencing is integral to immune tolerance of the placenta, but its suppression seems to be inconsistent across hTSC derivation methodologies.

Methylation-regulated targets in hESCs (Figure S4C) include two retrovirus-derived placental genes: *ERVW-1* (syncytin)^68^, which is critical in placental cell fusion and *ERVH48-1* (suppressyn), which protects placental cells from infection by Type D retroviruses^69^ (Figure 3J, 4K, S4D). We also observe upregulation of a set of genes that are typically expressed in placenta and germline but not elsewhere in development (*MAGEA4, MAGEA8, DDX43, BRDT, NAA11, TRIM60*) (Figure S4C). None of these genes have established functions in placenta, but the MAGE-A family of genes is widely activated in cancer and promotes cell growth by promoting ubiquitination and downregulation of p53^70^ and the nutrient-sensor AMPK^71^. In the murine germline, the *MAGE-A* genes have been established to suppress p53 protein levels and promote germ cell survival upon DNA damage or starvation^72^, which is intriguing in light of the weaker p53 response observed in hTSCs (Figure 3H, S3D and below).

An intriguing example featuring both placental and embryonic-specific methylation is the previously uncharacterized *PAX8* locus. PAX8 is a transcription factor implicated in kidney and thyroid development^73,74^ that is not expressed in placenta. PAX8 contains an internal antisense transcript called *PAX8-AS1*. In trophoblast, the *PAX8-AS1* promoter is unmethylated and highly expressed, and methylation is observed over the PAX8 promoter (Figure 4L). The phenomenon of transcriptional readthrough inducing methylation of an adjacent promoter has been observed in other contexts, notably imprinted genes^75,76^, and the function of *PAX8-AS1* may be to promote *PAX8* methylation and block its expression in placenta. In hESCs by contrast, *PAX8-AS1* is repressed by methylation, and the *PAX8* promoter is unmethylated and thus retains capacity for expression in subsequent development.

In total, while DNA methylation is not a general mechanism for lineage-specific silencing in epiblast or trophoblast, there are key examples of this phenomenon in each cell type.

### Trophoblast stem cells are more tolerant to DNA damage than embryonic stem cells

As shown above, deletion of DNMT1 causes hESCs, but not hTSCs, to upregulate p53 targets and be rapidly depleted from culture in a p53-dependent manner (Figures 3B, 3G-I, S3C). Positive signal for cleaved-caspase 3 was rapidly observed following *DNMT1* ablation in hESCs, but not in hTSCs, suggesting that only hESCs are initiating apoptosis in response to DNA hypomethylation (Figure S5A,B). It is well established in other cell types that DNA hypomethylation can cause mitotic instability, DNA damage and apoptosis^23,77^. We sought to determine whether DNMT1 loss causes DNA damage in hTSCs and hESCs, and whether differences in p53-dependent responses could explain why hTSCs can tolerate DNMT1 loss.

Loss of DNMT1 did not affect the percentage of hTSCs and hESCs positive for proliferation marker Ki67 or cause widespread enrichment for γH2AX, a marker for DNA double-stand breaks, in hTSCs (Figure S5C-E). However, we noted an increase in the number of polyploid cells in d7 and d10 DNMT1 KO hTSC (Figure S5F). This suggests that DNA hypomethylation may lead to mitotic abnormality and aneuploidy. DNMT1 KO hTSCs showed a higher frequency of multipolar spindles with supernumerary centrosomes during mitosis compared to *DNMT1^+/+^*hTSCs (Figure 5A,B). Chromosomal spreads of *DNMT1^-/-^* hTSCs further revealed a greater incidence of aneuploidy, with DNMT1 KO hTSCs frequently losing one or more chromosomes (Figure 5C,D, S5G). D7 DNMT1 KO *TP53^-/-^* hESCs similarly showed aneuploidy, with most affected cells exhibiting chromosome loss (Figure 5C,E & S5H). Collectively, these data show that DNMT1 loss causes mitotic abnormality in both hESC and hTSC, an effect masked in hESCs by rapid p53-induced lethality.

**Figure 5.**
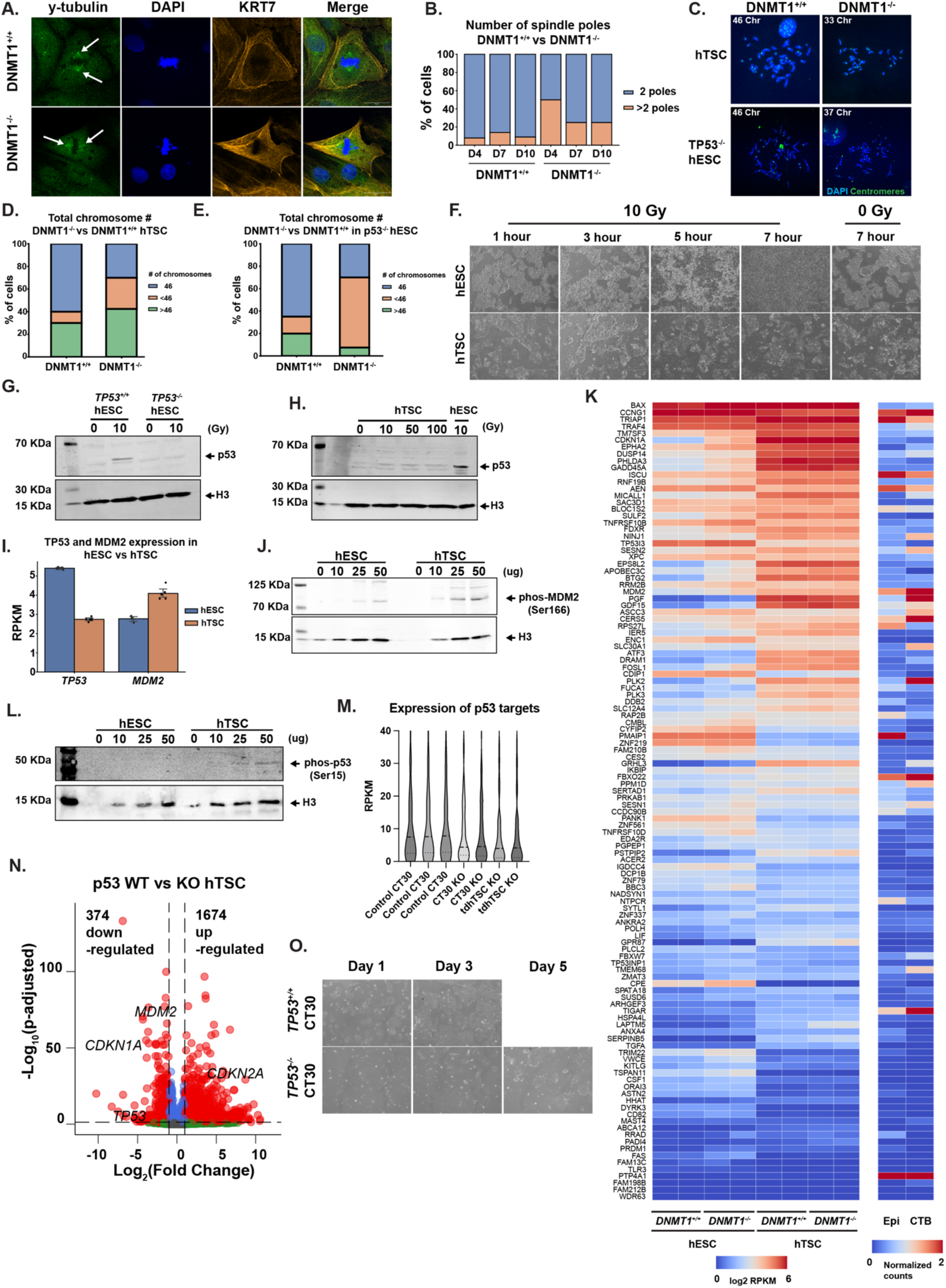
hTSCs have a diminished response to DNA damage despite active p53. (A) Immunofluorescence staining of centrosome poles during metaphase. hTSCs were stained for ã-tubulin (Green), KRT7 (Orange) and DAPI (Blue) at 100x magnification. Scale bar measures 25 μm. Arrows show enrichment of γ-tubulin demarking the centrosome. (B) Quantification of spindle poles in both *DNMT1^+/+^*and *DNMT1^-/-^* hTSCs from pictures in (A). (C) Immunofluorescence staining of day 7 *DNMT1^+/+^*and *DNMT1^-/-^* hTSCs and *TP53^-/-^* hESCs chromosomal spreads. Cells were stained with DAPI and a Human pan-centromeric DNA FISH probe (Green) at 100x magnification. (D-E) Quantification of chromosome numbers from immunofluorescent pictures in C). (F) Timelapse brightfield images of wild-type hESCs and hTSCs exposed to either 0 Grays or 10 Grays of ionizing radiation. Pictures were taken at 4x magnification. Scale bar measures 750 μm. (G) Western blot images of p53 and H3 enrichment in *TP53^+/+^* and *TP53^-/-^* hESCs after 0 Grays or 10 Grays of ionizing radiation exposure. Samples were collected three hours after exposure. (H) Western blot image showing p53 and H3 enrichment in hTSCs exposed 0, 10, 50 and 100 Grays of ionizing radiation. A wild-type hESCs positive control was included in the final lane. (I) Quantification of normalized read counts of wild-type hESC (blue) and hTSC (orange) for expression of *TP53* and *MDM2*. (J) Western blot image for phosphorylated MDM2 (Serine 166) and H3 enrichment in hESCs and hTSCs. For each sample, increasing input of protein lysate ranged from 5μg to 50 μg. (K) Quantification of normalized reads of downstream p53 targets genes in *DNMT1^+/+^* and *DNMT1^-/-^*hESCs and hTSCs, as well as average normalized read counts from single-cell analysis of Epiblast and cytotrophoblast cells in human embryos. (L) Western blot analysis of phosphorylated p53 (serine 15) and H3 in hESCs and hTSCs samples. For each sample, increasing input of protein lysate ranged 5 μg to 50 μg. (M) Violin plot of normalized read count of downstream p53 target genes (gene list from (K)) in wild-type hTSCs, *TP53^-/-^* hTSCs and *TP53^-/-^* transdiff hTSCs (tdhTSCs). (N) Volcano plot of differentially expressed genes comparing *TP53^+/+^* and *TP53^-/-^* hTSCs. Significant differentially expressed genes (fold change > 2 and p-adjusted value < 0.05) are marked in red. (O) Timelapse brightfield pictures comparing *TP53^+/+^* and *TP53^-/-^* hTSCs growth rates. 50K cells were seeded on day 0 and picture were taken on day 1, 3 and 5.

We then sought to understand whether p53 was inactive in the hTSCs relative to hESCs. To examine p53 activity in hESCs and hTSCs, we exposed both cell types to 10 Grays (Gy) of ionizing radiation. hESCs showed rapid cell loss after three hours post-ionizing radiation exposure, whereas hTSCs survived and even continued to proliferate under the same ionizing radiation conditions (Figure 5F, Figure S5J). hTSCs responded to ionizing radiation with increased enrichment of γH2A.X, indicating that hTSCs still undergo and detect DNA damage (Figure S5I), they simply do not die in response. These distinct cellular responses to ionizing radiation were accompanied by differences in p53 stabilization. While p53 protein accumulated in hESCs following 10 Gy exposure (Figure 5G), p53 accumulation and rapid lethality were not observed in hTSCs even after exposure to 100 Gy (Figure 5H, Figure S5K), an extremely high dose of ionizing radiation^78^.

We considered the possibility that p53 response may generally be impaired in hTSCs. Indeed, hTSCs show lower p53 expression and high RNA and protein expression of the negative regulator of p53, MDM2, compared to hESCs (Figure 5I, J). Inhibition of MDM2 activity by Nutlin-3a, led to increased accumulation of p53 signal in hTSCs, indicating that high MDM2 expression suppresses p53 accumulation in hTSCs (Figure S5L). Treatment with the sarco/endoplasmic reticulum Ca2+ ATPase inhibitor thapsigargin^79,80^, which induces ER stress, could induce apoptosis in hTSCs (Figure S5M-O). Thus, hTSCs are capable of undergoing apoptosis but do not do so in response to mitotic abnormality or DNA damage.

Despite lower TP53 transcript expression, hTSCs show higher constitutive levels of activated^81^ (Ser15p) p53 (Figure 5L), as well as higher constitutive expression of canonical p53 target genes, such as *CDKN1A*/p21 compared to hESCs (Figure 5K & Figure S5P). We generated *TP53*^-/-^ hTSCs and transdifferentiated *TP53^-/-^* hESCs to trophoblast (tdhTSC), and found that these showed lower levels of *CDKN1A* and TP53 target genes (Figure 5M, N). Surprisingly, we observed slower cell division of the *TP53^-/-^* hTSC (Figure 5O). We observe higher expression of *CDKN2A* in *TP53^-/-^* hTSCs, a compensatory mechanism that has been observed in other cell types upon *TP53* loss^82^ (Figure 5N). Thus, while hTSCs do not activate p53 in response to aneuploidy or DNA damage, they are capable of undergoing apoptosis and indeed have some constitutive p53 activity. These lineage-specific differences in p53 regulation highlight the differential tolerance to DNA hypomethylation between embryonic and placental stem cell lines.

### The p53 mitotic surveillance system is activated during DNA hypomethylation in hESCs

The mechanism underlying p53 activation in hESCs upon DNA hypomethylation remains unclear. We noted that both *DNMT1^-/-^* hTSCs, and *TP53^-/-^ DNMT1^-/-^* hESCs, show a high frequency of mitotic errors and aneuploidy, and that such defects can lengthen mitosis^83^. Prolonged mitosis in turn activates the mitotic surveillance pathway, a p53-dependent quality control mechanism, leading to G1/S cell-cycle arrest or apoptosis in the daughter cells as a safeguard for genomic integrity^84,85^. We hypothesized that *DNMT1^-/-^* hESCs may activate the p53-dependent mitotic surveillance pathway, rapidly clearing the abnormal cells, while this pathway is impaired in hTSCs. Analysis of genes involved in the mitotic surveillance pathway revealed reduced levels of *TP53BP1* transcript and protein in hTSCs relative to hESC (Figure 6A-C & Figure S6A), a key component of the mitotic surveillance pathway. We treated both hESCs and hTSCs with centrinone, a Polo-like Kinase 4 (PLK4) inhibitor that activates the p53 mitotic surveillance pathway by disrupting centriole duplication and extending mitosis^86^. hESCs undergo cell loss within 24 hours of centrinone treatment and upregulate p21 expression. In contrast, hTSCs tolerate centrinone treatment and continue to proliferate, albeit at a slower rate, with p21 already elevated under control conditions and unresponsive to centrinone (Figure 6D,E & Figure S6B).

**Figure 6.**
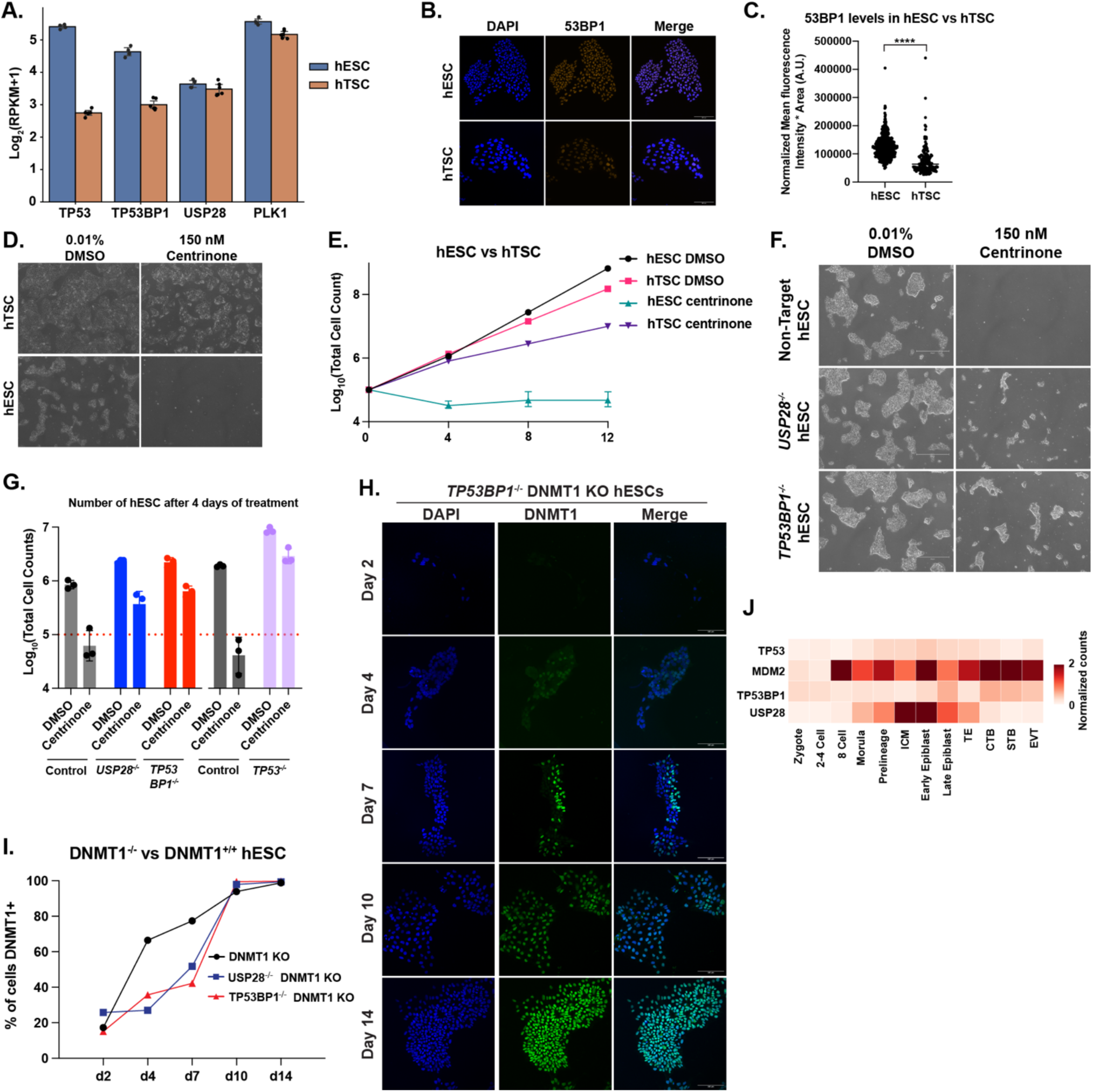
The mitotic surveillance pathway is activated in DNMT1 knockout cells. (A) Quantification of relative expression of key genes in the mitotic surveillance pathway hESCs (blue) and hTSCs (orange) from RNAseq samples. (B) Immunofluorescence staining of 53BP1 in hESCs and hTSCs. Cells were stained for 53BP1 (Orange), and DAPI (Blue) at 20x magnification. Scale bar measures 100 μm. (C) Quantification of 53BP1 protein signal per nuclei of imaged cells from (B). 53BP1 signal was normalized by H3 signal (not shown) ****: p< 0.0001 (D) Brightfield pictures of hESCs and hTSCs exposed to 150 nM centrinone or 0.01% DMSO at 4x magnification. (E) Quantification of cell growth in hESCs and hTSCs when treated with DMSO or 150 nM centrinone over the course of 12 days. Note the tolerance of hTSCs to centrinone treatment. (F) Brightfield pictures of Non-Target (control), *USP28^-/-^*, *TP53BP1^-/-^* hESCs treated with DMSO or centrinone over the course of 4 days. Pictures taken at 4x magnification. Scale bar shows 750 μm. (G) Quantification of Non-Target, *USP28^-/-^*, *TP53BP1^-/-^* and *TP53^-/-^* hESCs treated with centrinone or DMSO over the course of four days. Red dashed line indicates number of cells plated. *TP53^-/-^* and corresponding control were conducted at a separate time and are shown as a distinct control. (H) Immunofluorescent staining for DNMT1 in a bulk population of *TP53BP1^-/-^* hESCs 2 to 14 days after nucleofection with CRISPR/Cas9 and sgRNA targeting DNMT1. Cells were stained for DNMT1 (green) and DAPI (blue). Pictures were taken at 20x magnification. Scale bar measures 100 μm. (I) Percentage of Non-Target, *USP28^-/-^*, and *TP53BP1^-/-^* hESCs positive for DNMT1 after nucleofection with CRISPR/Cas9 and sgRNA targeting DNMT1 (DNMT1 KO). (J) Single cell analysis of TP53, MDM2, USP28 and TP53BP1 expression levels in human embryos from Zygote to Epiblast and placental cells.

To determine whether DNMT1 loss led to activation of the mitotic surveillance pathway, we knocked out the surveillance pathway genes *USP28* and *TP53BP1* hESCs (Figure S6C-E). *USP28^-/-^*, *TP53BP1^-/-^* and *TP53^-/-^* hESCs all show increased tolerance of centrinone treatment compared to non-target controls, showing continued cell growth over the course of 4 days (Figure 6F,G & Figure S6F). Only *TP53^-/-^*hESCs had increased resistance to lethality from ionizing radiation (Figure S6G), consistent with the *USP28^-/-^* and *TP53BP1^-/-^* showing a specific defect in the mitotic surveillance pathway rather than a general increase in tolerance for DNA damage. Both *USP28^-/-^* and *TP53BP1^-/-^* showed increased viability after DNMT1 KO, similar to *TP53^-/-^* hESCs (Figure 6H,I and Figure S6H). Functionally analogous to what we observed in hESC and hTSC, single-cell RNA-seq data from human embryos^65^ indicate that primed epiblast expresses lower levels of *MDM2* and higher levels of *USP28* than trophectoderm and CTBs (Figure 6J & Figure S6I-M).

Collectively, our findings are consistent with a model by which DNA hypomethylation leads to activation of p53 through the mitotic surveillance pathway in hESCs, preventing the survival of karyotypically abnormal cells. The mitotic surveillance system is defective in hTSCs, mimicking the *USP28^-/-^* or *TP53BP1^-/-^* hESCs and resulting in the accumulation of cells with chromosomal abnormalities.

## DISCUSSION

TRIM28 is clearly critical for trophoblast viability, as *TRIM28^-/-^*hTSCs do not survive in extended culture. This is especially intriguing considering that both naïve and primed hESCs can tolerate the loss of TRIM28 with no major defects in cell proliferation or viability^51^. The lethality of *Trim28*-deficient mice and mESC is accompanied by dramatic upregulation of transposons, including ∼100-fold upregulation of highly mutagenic IAP-Ez elements^8^, to the extent that transcriptional condensates are diverted away from pluripotency-associated superenhancers and toward ERV loci^87^. It should be noted here that mice contain a more aggressive suite of transposons, with transposable elements responsible for approximately 10% of mutations in mice but only 0.3% in humans^88,89^. Transposons may be likewise responsible for lethality in *TRIM28^-/-^*hTSCs, but no comparable upregulation of transposons is observed in TRIM28-deficient human cells. Instead, TRIM28 apparently suppresses the enhancer activity of LTRs, and the observed loss of *TRIM28^-/-^* hTSCs may be caused by aberrant expression of any one of hundreds of genes upregulated upon its loss.

A massively parallel reporter assay recently demonstrated enhancer activity for many individual MER11 elements^90^. We observe disparate activities of different subclasses of MER11A elements, with some showing evidence of constitutive enhancer activity in trophoblast and others gaining putative enhancer activity only upon TRIM28 loss. MER11A elements contain binding sites for transcription factors shared in placenta and a variety of primarily mesendodermal cell types, and differing levels of individual KRAB-ZFNs might regulate which MER11As show enhancer activity in which tissue. Consistent with this possibility, ZFN808 suppresses liver cell fate in pancreatic tissue by suppressing MER11A elements, and humans deficient for *ZNF808* show pancreatic agenesis^91^. It may also be the case that a sufficient dose of transcription factor can overwhelm TRIM28’s silencing capacity: the most active LTR7 sites in hESCs show as much TRIM28 binding as less active sites, but they have evolved to feature greater binding by pioneering KLF4 and SOX2 transcription factors^92^. Likewise, the high levels of GATA3 and TFAP2C in trophoblasts may overcome TRIM28-mediated silencing that is effective in other cell types.

DNA methylation eventually supplants TRIM28 as a primary mediator of transposon repression. This likely occurs by multiple mechanisms. The suppression of transcription and H3K4 methylation at active sites facilitates methylation by the *de novo* methyltransferases DNMT3A and DNMT3B^93^. Furthermore, the maintenance methyltransferase DNMT1 has apparent *de novo* activity at TRIM28 sites, either via direct recruitment or because H3K9 methylation stabilizes UHRF1 binding and recruits DNMT1^94^.

The role of DNA methylation in lineage-specific gene regulation has been a point of some contention^95^. Methylation is clearly critical in the regulation of imprinted and germline genes, but this methylation is established in gametogenesis or early development respectively and is essentially ubiquitous and static throughout the soma. Because epiblast and trophoblast cells diverge prior to the wave of *de novo* methylation upon implantation, somatic and placental tissue have dramatically different methylomes, making lineage-specific regulation more plausible in this context. We were able to identify a small set of genes repressed by DNA methylation selectively in hTSC, with the strongest examples showing corresponding transcriptional increases in CTB treated with a DNMT1 inhibitor. We also identified a larger set of genes repressed by DNA methylation in hESCs. Interestingly, there was no obvious pattern in function, etiology, gene structure or evolutionarily age of methylation-repressed target genes. *CDX1*, *CRABP1*, *NNAT*, *DPPA4* and *HLA-A* have essentially nothing in common beyond needing to not be expressed in trophoblast. We observed instances of nested genes (*NNAT, PAX8-AS1*), and virus-derived genes (*ERVW-1*, *ERVH48-1*) regulated by methylation but these did not reflect any general case. Lineage-specific methylation apparently arises sporadically during evolution and becomes critical for the regulation of rare, individual genes.

Loss of DNMT1 facilitates conversion of murine embryonic stem cells to trophoblast^96^, leading to the hypothesis that methylation of “gatekeeper genes” in embryonic lineage, including core placental transcription factors such as *Elf5*, blocks conversion to trophoblast^97^. Our data do not contradict this possibility. We only identify genes that are upregulated upon DNMT1 loss in steady state culture conditions; a gene whose activation requires both demethylation and additional changes in cellular signaling will not be observed. Thus, we have found a minimal rather than maximal set of methylation-regulated genes.

We and others find that loss of DNA methylation led to mitotic defects and karyotypic abnormalities^23,77^. Three possible mechanisms by which DNA hypomethylation could contribute to karyotypic abnormality include 1) de-repression of centromeric and peri-centromeric regions, 2) activation of meiosis genes, and 3) activation of transposable elements. The centromere consists of α-satellite monomer repeat elements, while the flanking peri-centromeric regions are more heterogeneous in nature with α-satellite monomer and other transposable elements such as SINEs and LINEs. DNMT3s are recruited to centromeric protein complexes and deposit DNA methylation over these regions^98–100^. Directed hypomethylation of centromeres results in centromeric structural aberrations and aneuploidy^101^. We observe increased expression of satellite RNA upon DNMT1 loss, and satellite RNA is known to interfere with proper kinetochore assembly at least in the context of meiosis^102^. DNA hypomethylation also leads to the activation of meiotic genes in hESCs and hTSCs. Expression of meiotic genes can interfere with proper mitosis and ectopic expression of meiotic genes in cancer leads to chromosomal instability and aneuploidy^103,104^. Thus, it is possible that *DNMT1^-/-^* hESCs or hTSCs undergo defective mitosis due to ectopic meiotic gene expression. Finally, although there is no direct proof that transposon expression can cause pervasive karyotypic abnormality, it is established that transposition causes DNA double-strand breaks and *mael-*null oocytes, or oocytes overexpressing LINE elements, show increased rates of karyotypic abnormality during meiosis^105^. Which of these mechanisms drives karyotypic abnormality in *DNMT1*-null cells is unclear, but regardless, this abnormality then induces fast mitotic surveillance pathway-mediated lethality in hESCs and slower lethality in hTSCs.

There is extensive evidence that the human placenta is generally more likely to contain mutations and mosaic karyotypic abnormalities than fetal tissue^106–109^. We find that hTSCs have a reduced general propensity for apoptosis and remarkable tolerance for DNA damage, which may explain why mutations and karyotypic abnormalities are so common. Consistent with our data, murine embryos induced to undergo aneuploidy show extensive apoptosis of aneuploid cells in the inner cell mass but far less in the trophoblast^110^. Across metazoans, it has been observed that polyploid cells have a dampened DNA damage response, although not by any consistent mechanism across cell types^111^. hTSCs show some propensity for polyploidy^24^, and endoreplication and fusion are established phenomena in EVT and STB differentiation respectively. In addition to higher *MAGE* gene expression and lower *TP53BP1*, hTSCs express much higher levels of *11NTP63α*, a *TP63* isoform which antagonizes canonical p53 pro-apoptotic and anti-proliferative signaling activity^112,113^. Notably, *11NTP63α* expression has been linked to radio-resistance in various squamous carcinoma cell lines^113–115^.

The trophoblast is an anomalous cell that had to evolve a specialized function very quickly. In doing so, it has co-opted transposons to regulate genes, used methylation to silence isolated but important loci, and been forced to tolerate a high degree of karyotypic abnormality as the price of its unique properties.

## RESOURCE AVAILABILITY

### Lead contact

Requests for cells, materials, or additional information can be sent to William A. Pastor.

### Materials availability

Cell lines and plasmids generated in this project will be made readily available by the lead contact.

### Data availability

At present, we cannot upload primary sequencing data from this study to the Gene Expression Omnibus (GEO) database because of the ongoing government shutdown. TRIM28 ChIP-seq data can be visualized via this UCSC genome browser session: https://genome.ucsc.edu/s/DSaini/TRIM28_ChIPseq_hTSC Primary sequencing data will be made available to interested parties upon request.

## ACKNOWLEDGEMENTS

We thank Nobuko Yamanaka for teaching us how to implement karyotyping assays. We thank the Centre for Applied Genomics at SickKids Hospital, Canada’s Michael Smith Genome Sciences Centre at BC Cancer, the Goodman Cancer Institute Flow Cytometry core, and the McGill University Advanced BioImaging Facility (ABIF) for dedicated service. The research was funded by New Frontiers in Research Fund (NFRF) grant NFRFE-2018-00883 to W.A.P. and R.S., CIHR Project Grant PJT-166169 and NSERC Grant RGPIN-2018-04856 to W.A.P, Wellcome Trust Career Development Award 301798/Z/23/Z to J.M.F, NIH grants HD113673, HD119510, HD103161, HD062546, HD101319 to S.P., Réseau de Recherche sur le Cancer of the Fonds de Recherche du Québec-Santé, Canada Foundation for Innovation (CFI) 33902, CIHR FDN-143281 and Cancer Research Society grant 263225 to M.P, an FRQS Junior Scholar 1 award and CIHR PJT-198136 to R.C.-M. D.S. was supported by an FRQ-NT Graduate fellowship and a Carole Epstein fellowship in Women’s Health. W.A.P. is an FRQS Chercheur-boursier.

## AUTHOR CONTRIBUTIONS

D.S., M.S.K., S.G.A.B., T.E.I., S.R., A.N., J.K.C., S.D. and T.S. performed experiments. D.S., M.S.K., S.C.V., A.F and R.M. performed bioinformatic analysis. R.S., M.P., J.M.F., S.P, R.C.-M. and W.A.P. supervised research. D.S. and W.A.P. wrote the manuscript.

---

**Figure S1.**
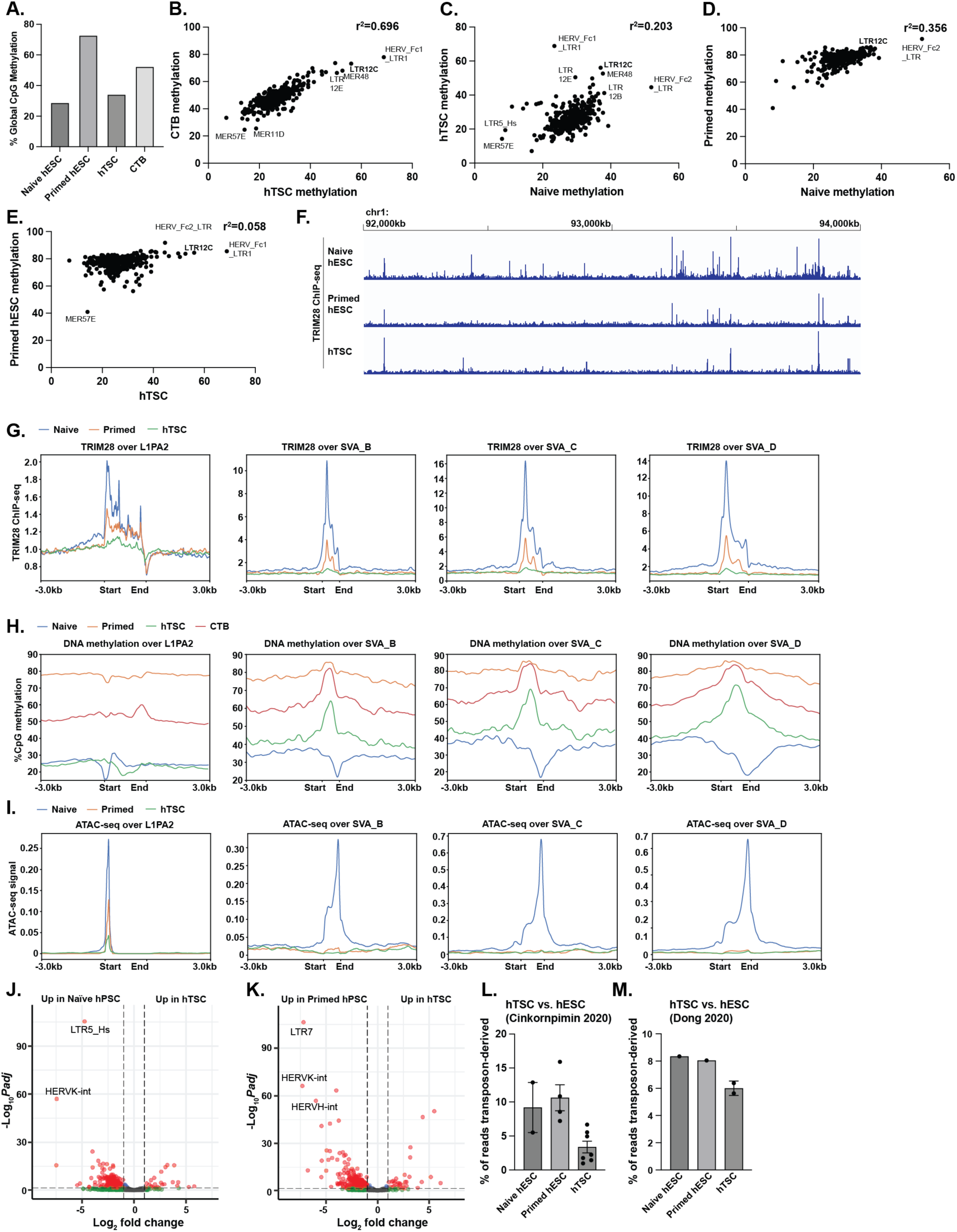
Distribution of TRIM28 and 5mC over transposons and transposon expression in in embryonic and trophoblast stem cells, related to Figure 1. (A) Global %CpG methylation of cell types indicated. (B - E) Average CpG methylation level of each LTR class in cell types indicated, with correlation coefficient calculated. Note high correlation of LTR methylation between CTB and hTSC. (F) TRIM28 ChIP-seq signal over a 2MB region on chromosome 1. Note the presence of far more TRIM28 sites in Naïve hESC relative to Primed or hTSC. (G) Metaplot of TRIM28 ChIP-seq signal over transposon classes indicated in naïve hESC, primed hESCs, and hTSCs. (H) Metaplot of %CpG methylation over transposon classes indicated in naïve hESC, primed hESCs, and hTSCs. (I) Metaplot of ATAC-seq signal over transposon classes indicated in naïve hESC, primed hESCs, and hTSCs. (J) Volcano plot of expression of transposon classes in hTSC vs. naïve hESC. (K) Volcano plot of expression of transposon classes in hTSC vs. primed hESC. (L, M) Percentage of transposon-derived reads in cell type indicated for data source indicated.

**Figure S2.**
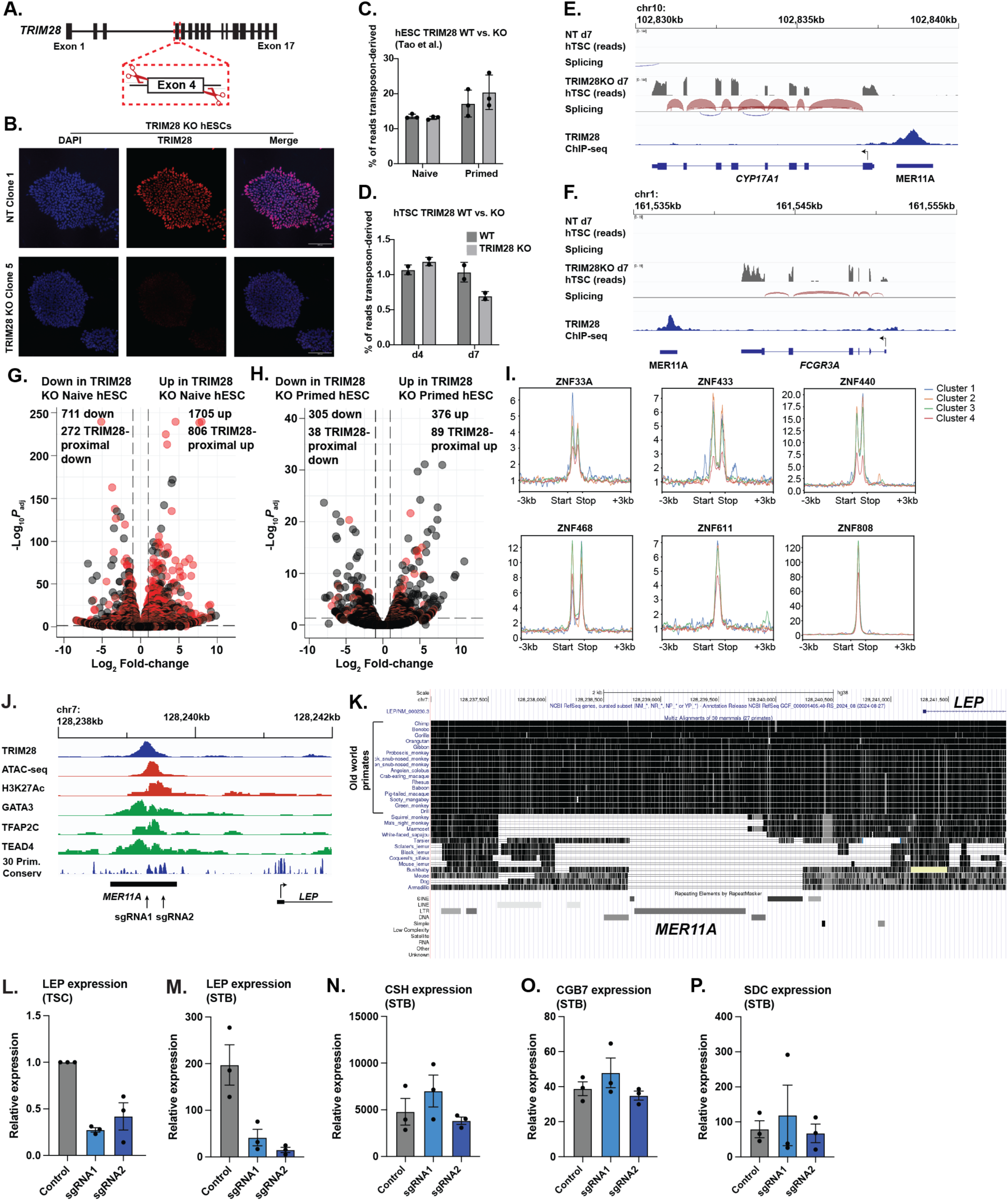
TRIM28 regulates adjacent genes by suppression of LTR transposons, related to Figure 2. (A) Schematic of targeting of *TRIM28* locus. (B) Immunofluorescent staining for TRIM28 in a TRIM28 KO hESC clonal line and a control clonal line nucleofected with TRIM28-targeting sgRNA or non-targeting sgRNA. hESC were stained with TRIM28 (red) and DAPI (blue). Pictures were taken at 20x and the scale bar measures 100 μm. (C) Percentage of transposon-derived reads from control and TRIM28^-/-^ naïve and primed hESCs. Data is derived from Tao et al. 2018. (D) Percentage of transposon-derived reads from control and bulk TRIM28 d4 and d7 KO hTSC. (E, F) RNA-seq and ChIP-seq plotted over the *CYP17A1* (E) and *FCGR3A* (F) loci in control and d7 *TRIM28* KO hTSCs. Note that while both genes are highly upregulated in the TRIM28 KO, there is no splicing of transcripts from the MER11A element to the genes. (G, H) Volcano plot of TRIM28 KO vs. control naïve (G) and primed (H) hESCs. TRIM28-proximal (<50kb distance) genes are indicated in red. (I) Metaplots of six KRAB-ZFN proteins over the four MER11A clusters identified in Figure 3F. Most show uniform enrichment over Clusters 1 - 3, but ZNF468 shows lower enrichment over the less heterochromatinized Cluster 1. Note that this ChIP-seq data is derived from ectopic expression of KRAB-ZFNs in 293T cells. (J) ATAC-seq and ChIP-seq enrichment plotted over a 4kb region upstream of and including the *LEP* promoter. Sites of two sgRNA used for CRISPRi targeting of the locus are indicated. (K) Conservation of region upstream of *LEP* in twenty-six primate and three non-primate mammals. Note that the MER11A element is only present in old world primates. (L, M) Expression of *LEP* in hTSCs and STBs expressing sgRNA targeting sites indicated in (J) or a non-targeting control sgRNA. Expression is normalized to level in control hTSC. (N – P) qPCR data showing expression of the STB markers *CSH* (N), *CGB7* (O) and *SDC* (P) in STBs expressing a CRISPRi system and the sgRNA indicated in (J). Note normal induction of these STB genes in all three samples, indicating the sgRNA had no general effect on STB differentiation.

**Figure S3.**
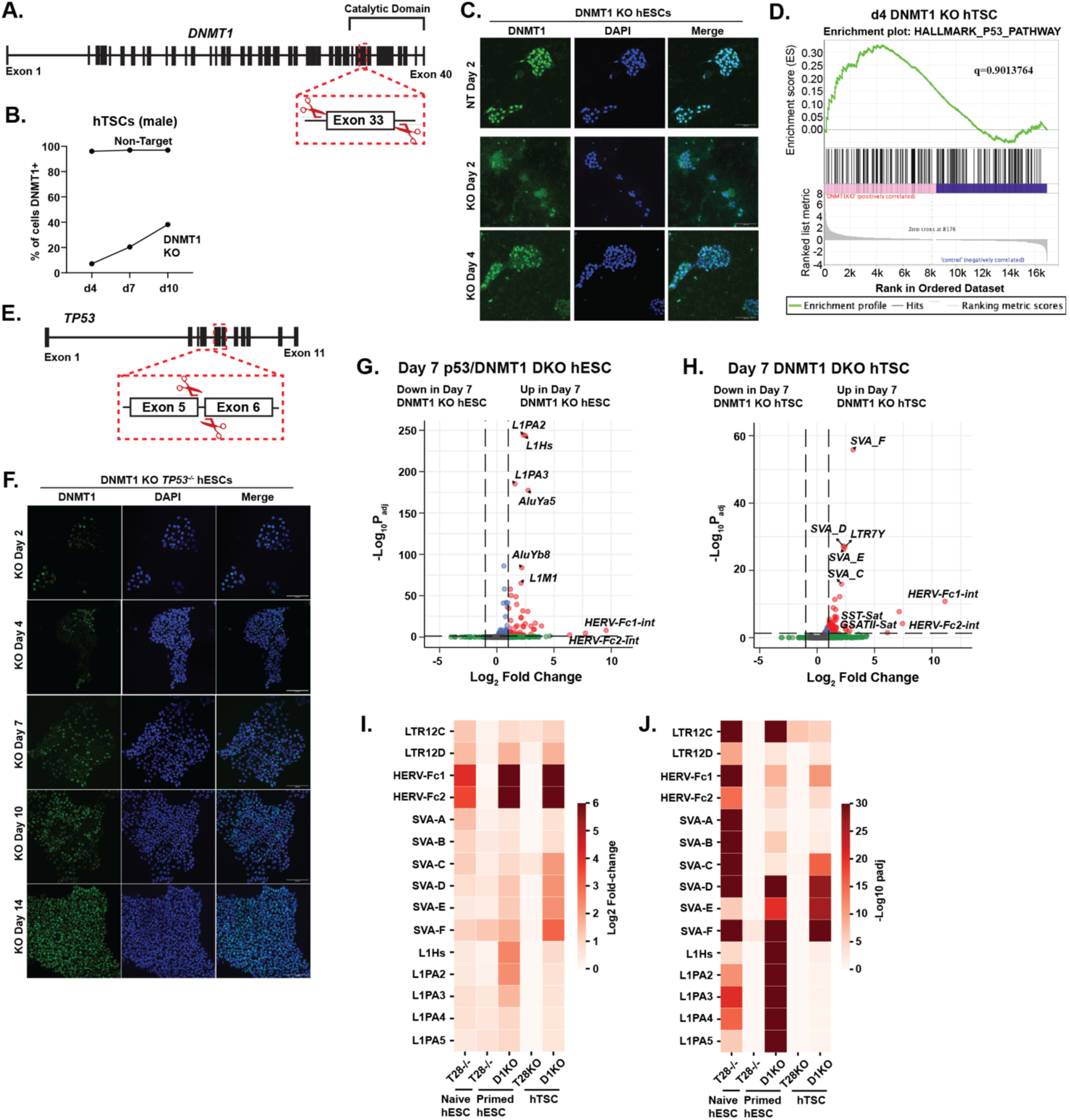
DNA methylation is essential for survival and transposon repression of hTSCs and hESCs, related to Figure 3. (A) Schematic of targeting of *DNMT1* locus. (B) Percentage of CT29 hTSC positive for DNMT1 after nucleofection with CRISPR/Cas9 and sgRNA targeting DNMT1 (DNMT1 KO) or non-targeting sgRNA (Non-target). Rate of loss for the male CT29 line is similar to the rate of loss for the female CT30 line as shown in Figure 3A. (C) Immunofluorescent staining for DNMT1 in a bulk population of hESCs 2 or 4 days after nucleofection with CRISPR/Cas9 and sgRNA targeting DNMT1. hESCs that underwent nucleofection with a non-targeting sgRNA are shown as a control. Note rapid loss of *DNMT1* deficient hESCs. hESCs were stained for DNMT1 (green) and DAPI at a magnification of 20x and the scale bar measures 100 μm. (D) GSEA analysis comparing expression of P53 pathway genes in day 4 DNMT1 KO hTSCs and controls. No statistically significant difference increase is observed in the DNMT1 KO cells. (E) Schematic of targeting of *TP53* locus. (F) Immunofluorescent staining for DNMT1 in a bulk population of *TP53^-/-^* hESCs after nucleofection with CRISPR/Cas9 and sgRNA targeting DNMT1. hESC’s that underwent nucleofection with a non-targeting sgRNA are shown as a control. Note slower loss of *TP53^-/-^* DNMT1 KO hESCs as compared with S2C. hESCs were stained for DNMT1 (green) and DAPI at a magnification of 20x and the scale bar measures 100 μm. (G, H) Volcano plot of transposon classes upregulated in DNMT1 KO hESC (G) and hTSC (H). (I, J) Fold change (I) and statistical significance of upregulation (J) for transposon class and cell type indicated. Note that elements such as SVAs and LINEs switch from negative regulation by TRIM28 in naïve hESCs to negative regulation by DNA methylation in primed hESCs and hTSC, in concert with the DNA methylation gain observed in Figure 1F-H and S1G-1.

**Figure S4.**
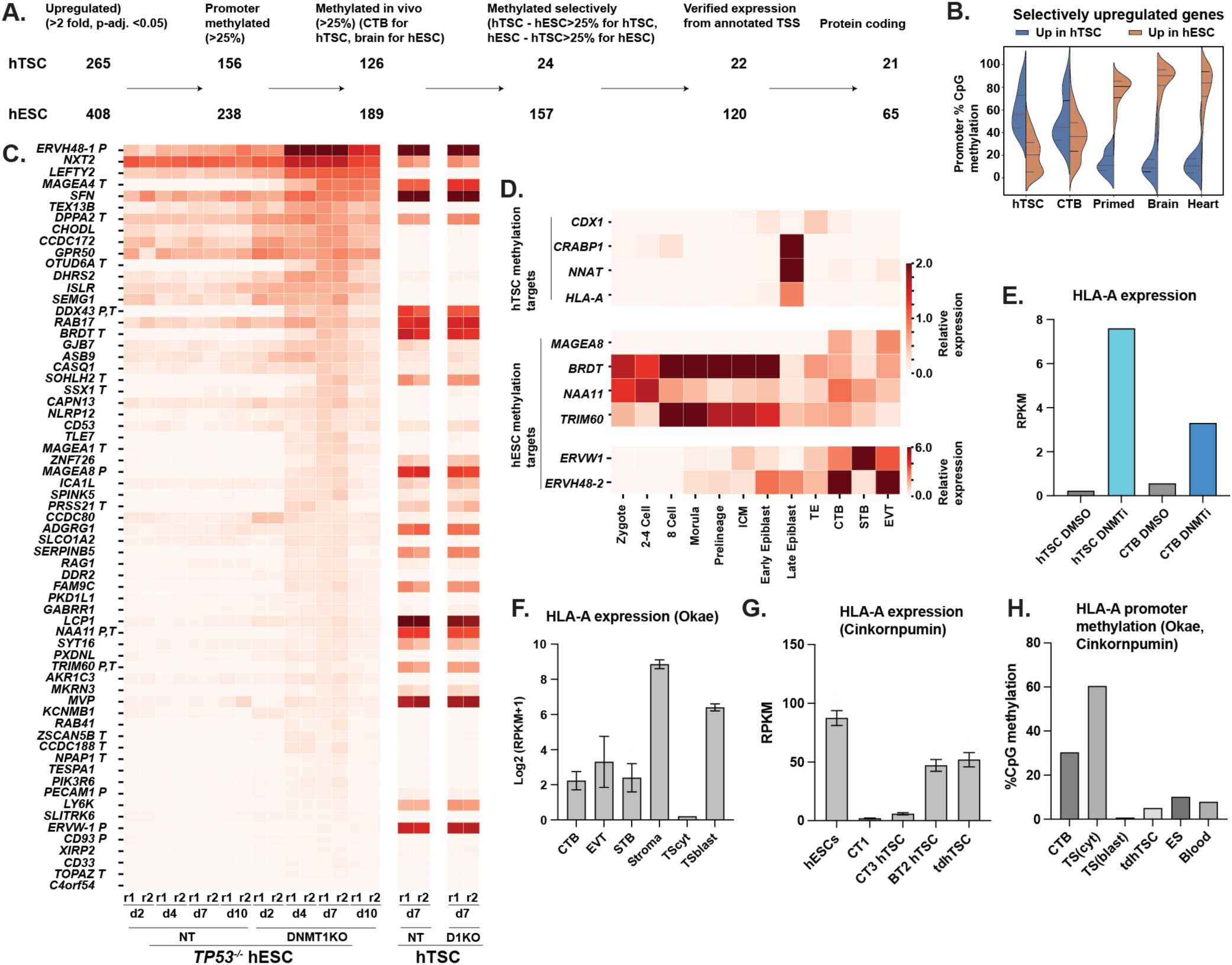
Regulation of developmentally important genes by DNA methylation in trophoblast and epiblast lineage, related to Figure 4. (A) Schematic for identification of genes regulated by 5mC selectively in one lineage. Starting from the genes only upregulated in one set in (A), required >25% promoter methylation (promoter: -500bp to 300bp), >25% methylation in placental or somatic tissue, methylated selectively in one tissue but not both. We next confirmed expression from the indicated promoter on Integrative Genome Viewer (IGV) to confirm a set of 5mC-regulated transcripts. The subset that are protein coding is finally indicated. (B) Split violin plot indicating promoter DNA methylation of the 22 hTSC-specific and 120 hESC-specific transcripts. (C) Heatmap showing expression of protein coding genes repressed by 5mC in hESCs in cell types indicated. Genes specifically expressed in placental (P) and/or testis (T) as determined by Human Protein Atlas are specifically labelled. (D) Single cell analysis of expression levels of representative hTSC and hESC methylation target genes in human embryos from zygote to epiblast and placental cells. (E) Expression of HLA-A in control and DNMT1i-treated hTSC and CTB. (F) HLA-A expression in tissues indicated, as well as placenta-derived (TScyt) and blastocyst-derived (TSblast) hTSC. (G) HLA-A expression in cell type indicated. tdhTSC refers refers to hTSC generated by transdifferentiation from hESC. (H) HLA-A promoter methylation in cell type indicated. All data are from Okae 2018, except for TS(transdiff) which is from Cinkornpumin 2020. Note concordance of hypomethylation and leaky expression of HLA-A in blastocyst-derived or transdifferentiated hTSC.

**Figure S5.**
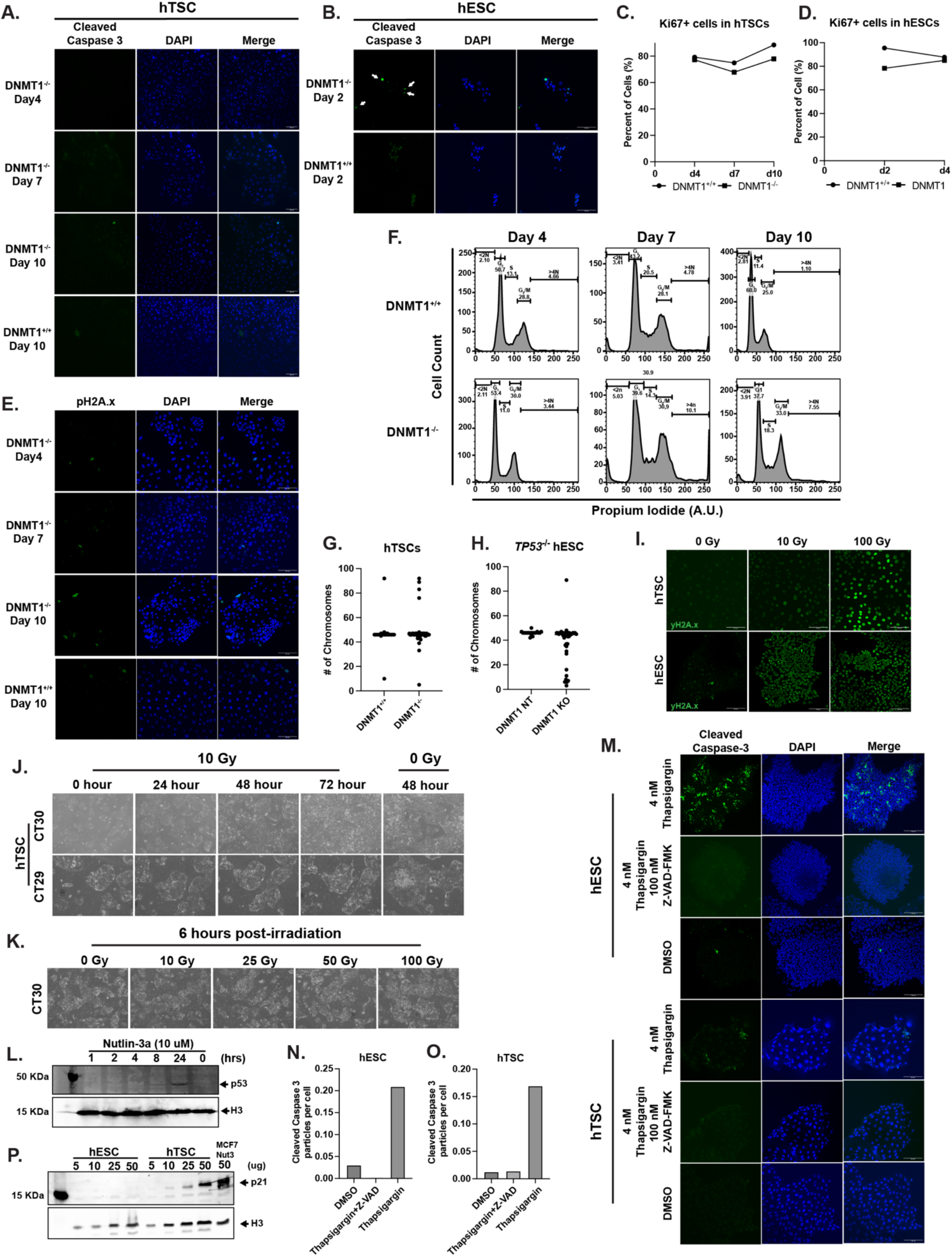
hTSCs have a diminished response to DNA damage despite active p53, related to Figure 5. (A,B) Immunofluorescent staining for active apoptosis in (A) hTSCs and (B) hESCs after DNMT1 ablation. *DNMT1^+/+^* and *DNMT1^-/-^* hTSCs were stained for cleaved caspase 3 (green) and DAPI (blue). Note the lack of active cleaved caspase 3 signal in *DNMT1^-/-^* hTSCs and arrows showing enrichment of positive cleaved caspase 3 in *DNMT1^-/-^* hESCs. Pictures were taken at 20x magnification and the scale bar measures 100 μm. (C,D) Quantification of Ki67 positive cells in *DNMT1^+/+^*and *DNMT1^-/-^* (C) hTSCs and (D) hESCs from immunofluorescent pictures. (E) Immunofluorescent staining of *DNMT1^+/+^* and *DNMT1^-/-^* hTSCs stained for yH2A.x (green) and DAPI (blue). Pictures were taken at 20x magnification and the scale bar measures 100 μm. (F) Flow cytometry of *DNMT1^+/+^* and *DNMT1^-/-^* hTSCs stained with propidium iodide to measure DNA content. Note the increase in polyploid cells in *DNMT1^-/-^* hTSCs in Day 7 and Day 10 samples. (G-H) Swarm plots quantification of chromosomal counts in Figure 5C. (I) Immunofluorescent staining of wild-type hTSCs and hESCs exposed to 0, 10 or 100 Grays of ionizing radiation. Cells were stained for yH2A.x (green). Note the increasing intensity of yH2A.x signal with increased exposure to ionizing radiation. Pictures were taken at 20x magnification and the scale bar measures 100 μm. (J) Timelapse brightfield picture of hTSCs, CT30 (female) and CT29 (male), exposed to 10 Grays of ionizing radiation. Pictures were taken every 24 hours to capture growth of cells after radiation exposure. Control hTSCs were not exposed to ionizing radiation (0 Grays) and seeded at the same density as radiation-treated cells. Pictures were taken at 4x magnification. (K) Brightfield pictures of hTSCs exposed to 0-100 Grays of ionizing radiation. Pictures were taken 6 hours post-radiation exposure. Note the tolerance of hTSCs to 100 Grays of ionizing radiation. Pictures were taken at 4x magnification and the scale bar measures 750 μm. (L) Western blot images of p53 enrichment in wild-type hTSC treated with 10 uM Nutlin-3a for 0 to 24 hours. 24 hours of Nutlin-3a treatment was required for positive p53 signal. (M) Immunofluorescent staining of hESCs and hTSCs exposed to either Thapsigargin, Thapsigargin and Z-VAD-FMK or DMSO. Cells were stained with cleaved caspase 3 (green) and DAPI (blue) to measure active apoptosis. Note enrichment of cleaved caspase 3 signal in Thapsigargin treated cells and the lack of signal when apoptosis is blocked with Z-VAD-FMK. Pictures were taken at 20x magnification and the scale bar measures 100 μm. (N, O) Quantification of images in M, showing cleaved-caspase3 particles per cell in (N) hESCs and (O) hTSCs. (P) Western blot analysis hESCs and hTSCs expression of p21 protein levels. Protein lysates were stained for both p21 and H3. Positive signal included MCF cells exposed to UV radiation. Increasing protein lysates from 5 to 50 ug were added per samples.

**Supplemental Figure 6.**
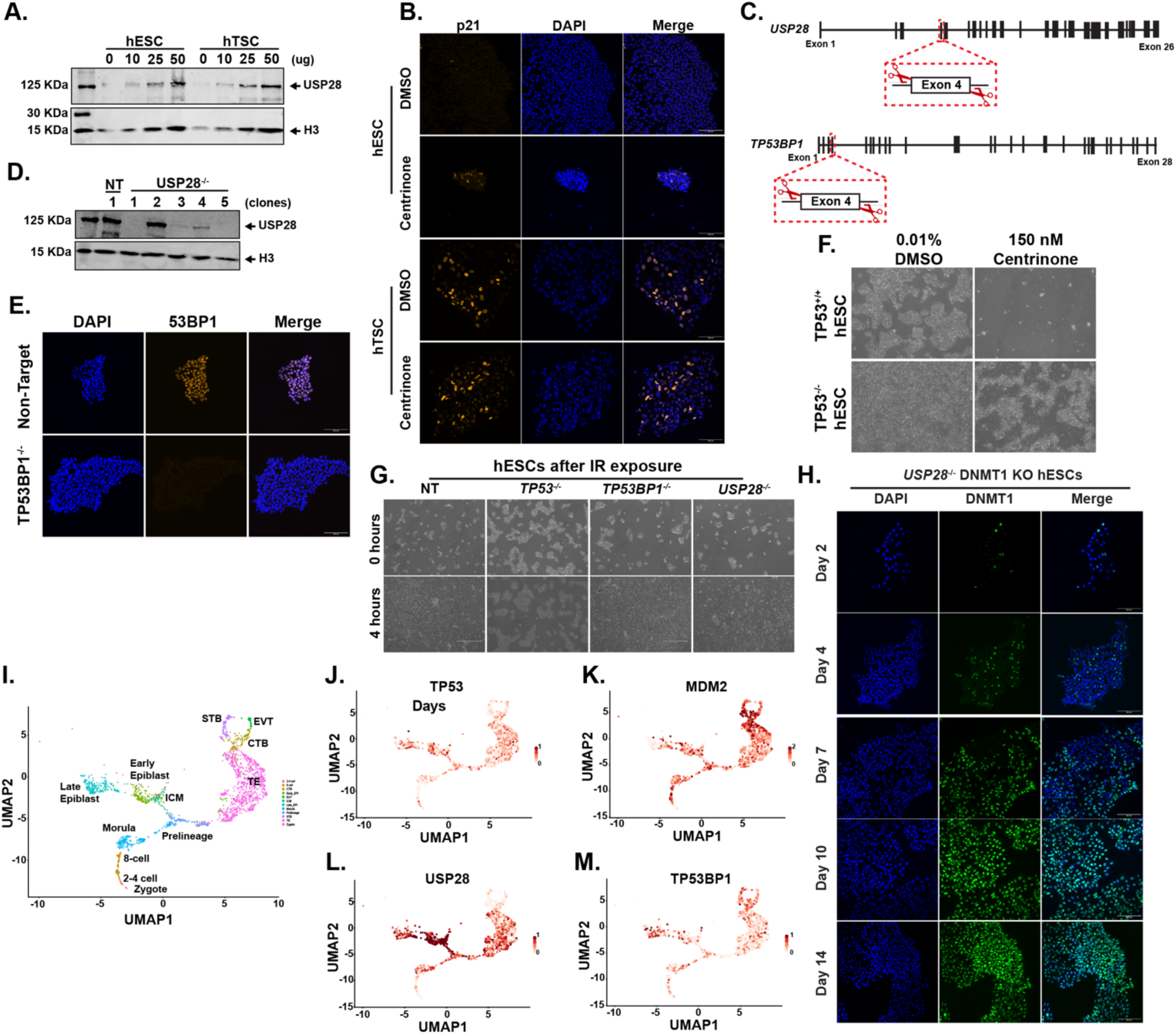
The mitotic surveillance pathway is activated in DNMT1 knockout cells, related to Figure 6. (A) Western blot analysis to USP28 protein level in hESCs and hTSCs. Protein lysates input was added in increased concentration from 5 ug to 50 ug. Lysates were stained for both USP28 and H3. (B) Immunofluorescent staining of mitotic surveillance activation after centrinone treatment in hESCs and hTSCs. Cells were treated with 150 nM centrinone or DMSO for 48 hours and stained for p21 (orange) and DAPI (blue). Pictures were taken at 20x magnification and the scale bar measures 100 μm. Note activation of p21 in hESCs exposed to centrinone and constant expression of p21 in hTSC. (C) Schematic of CRISPR-Cas9 strategy to target USP28 and TP53BP1 in hESCs. sgRNA guides (red scissors) were designed to flank Exon 4 to generate frameshift mutation. (D) Western blot analysis of hESC clones after USP28 ablation. Clones 1 and 5 were chosen for subsequent experiments. (E) Immunofluorescent staining of hESCs clones after 53BP1 ablation. hESCs were stained for 53BP1 (orange) and DAPI. Note lack of 53BP1 expression in the *TP53BP1^-/-^*hESCs. Pictures were taken at 20x magnification and the scale bar measures 100 μm. (F) Brightfield pictures of *TP53^+/+^* and *TP53^-/-^* hESCs after exposure to 150 nM centrinone. Cells were seeded at 100K in a 6-well plate and followed over the course of 4 days. Note tolerance of *TP53^-/-^* hESCs to centrinone treatment. Pictures were taken at 4x magnification. (G) Brightfield pictures of control, *TP53^-/-^*, *USP28^-/-^*, and *TP53BP1^-/-^* hESCs upon treatment with 10 Grays of ionizing radiation. Pictures were taken at 4x magnification and the scale bar measures 750 μm. (H) Immunofluorescent staining of *USP28^-/-^*hESCs after DNMT1 ablation. Cells were stained for DNMT1 (green) and DAPI (blue). Pictures were taken at 20x magnification and the scale bar measures 100 μm. (I) Re-analysis of single cell RNAseq from human embryos Seurat objects datasets from Zygote to Epiblast/CTB. (J-M) Expression heatmap profiles of *TP53, MDM2, USP28* and *TP53BP1* in during human embryo development.

