## Supplementary Table captions for "Divergent roles of DNA methylation, TRIM28, and p53 surveillance in human embryonic and trophoblast stem cells"

**Table S1.** Replicate information and mapping statistics for all next generation sequencing libraries generated in this manuscript.

**Table S2.** All datasets mined in this manuscript

**Table S3.** For each LTR class, the number of TRIM28 ChIP-seq peaks, the percent methylation, the percent methylation relative to surrounding sequence, and the ATAC-seq enrichment in the cell types indicated. Also shown is RAD analysis showing ratio of (upregulated in hTSC/upregulated in hESC) for genes within 50kb of an element of this class.

**Table S4** All differentially expressed gene calling and RPKM values for RNA-seq conducted in the course of this manuscript.

**Table S5** List of all TRIM28-bound LTR transposon within 50kb of a gene upregulated upon TRIM28 knockout in hTSCs.

**Table S6** Analysis to identify methylation-regulated genes, showing gene sets identified in Figure S4A.
